# Projecting ancient ancestry in modern-day Arabians and Iranians: a key role of the past exposed Arabo-Persian Gulf on human migrations

**DOI:** 10.1101/2021.02.24.432678

**Authors:** Joana C. Ferreira, Farida Alshamali, Francesco Montinaro, Bruno Cavadas, Antonio Torroni, Luisa Pereira, Alessandro Raveane, Veronica Fernandes

## Abstract

Arabian Peninsula is strategic for investigations centred on the structuring of the modern human population in the three main groups, in the awake of the out-of-Africa migration. Despite the poor climatic conditions for recovery of ancient DNA human evidence in Arabia, the availability of genomic data from neighbouring ancient specimens and of informative statistical tools allow better modelling the ancestry of these populations. We applied this approach to a dataset of 741,000 variants screened in 291 Arabians and 78 Iranians, and obtained insightful evidence. The west-east axis was a strong forcer of population structure in the Peninsula, and, more importantly, there were clear continuums throughout time linking west Arabia with Levant, and east Arabia with Iran and Caucasus. East Arabians also displayed the highest levels of the basal Eurasian lineage of all tested modern-day populations, a signal that was maintained even after correcting for possible bias due to recent sub-Saharan African input in their genomes. Not surprisingly, east Arabians were also the ones with higher similarity with Iberomaurusians, who were so far the best proxy for the basal Eurasians amongst the known ancient specimens. The basal Eurasian lineage is the signature of ancient non-Africans that diverged from the common European-East Asian pool before 50 thousand years ago, and before the later interbred with Neanderthals. Our results are strong evidence to include the exposed basin of the Arabo-Persian Gulf as possible home of basal Eurasians, to be investigated further on namely by searching ancient Arabian human specimens.

## Introduction

Coalescence analyses on modern-day genomic data have been basilar in the inference of the most important events along human evolution (Hellenthal, et al. 2014; Li, et al. 2008; Soares, et al. 2012). However, by 2010, the successful technical improvements on ancient DNA (aDNA) cataloguing began contributing essential insights into the overall picture (Skoglund and Mathieson 2018). In fact, the new possibility of surveying diversity that did not leave any genetic signature till the present time is revealing an extremely rich picture involving the interplay between several population groups. So, New insights have been obtained namely on the out-of-Africa migration (OOA; (Llorente, et al. 2015; Schuenemann, et al. 2017; Stringer 2014; Stringer and Andrews 1988)), the divergence between the non-African populations (Lazaridis 2018; Lipson and Reich 2017; Yang and Fu 2018), and their earliest movements into Eurasia and America (Fu, et al. 2014; Fu, et al. 2016; Lazaridis, et al. 2014; Posth, et al. 2018; Raghavan, et al. 2014; Rasmussen, et al. 2014). The aDNA catalogue already contains hundreds of whole-genome shotgun data and thousands of targeted capture data (SNP array-like) from specimens that date as far back as 45 thousand years ago (ka). Unfortunately, there is still a strong bias towards European, Siberian, Native American and Near Eastern specimens, so that large gaps remain in the reconstruction of Asian and African genomic histories (Pereira, et al. 2021; Skoglund and Mathieson 2018). A difficult-to-overcome factor responsible for this bias is the environmental influence in aDNA preservation, with arid deserts (AP-Arabian Peninsula, Sahara and other deserts) and humid tropical forests (large parts of Africa, Asia and America) amongst the worst possible conditions (Hambrecht and Rockman 2017). Nevertheless even for these regions, newly developed statistical tools are allowing to evaluate the contribution of neighbouring ancient founders to the genomes of current populations (Leslie, et al. 2015; Montinaro, et al. 2015; Raveane, et al. 2019).

AP played a major role or at least was in the path of the successful OOA migration at around 70-60 ka, and important genetic and archaeology findings have been uncovered in the region (Fernandes, et al. 2012; Groucutt, et al. 2018; Petraglia, et al. 2012; Rose, et al. 2011; Scerri, et al. 2018). Pleistocene Arabia was exposed to several climate change episodes that impacted modern human occupation (Petraglia, et al. 2020). During wet periods, populations would expand from the coast to hinterland following river valleys, while contracting to refugia during arid periods. Climatic and archaeological evidences indicate the existence of three main AP refugia (Rose 2010): the Red Sea coastal plain; the Dhofar Mountains and adjacent littoral zone in Yemen and Oman; and the exposed basin of the Arabo-Persian Gulf. Genomic studies conducted by us and others (Alshamali, et al. 2009; Cerný, et al. 2011; Cerný, et al. 2009; Fernandes, et al. 2012; Fernandes, et al. 2019; Fernandes, et al. 2015; Musilová, et al. 2011; Rodriguez-Flores, et al. 2016; Vyas, et al. 2017) in extant AP populations revealed a significant heterogeneity between western and eastern AP populations, with the former being like a continuum with sub-Saharan Africa and Levant, while the latter is more similar to Iranian and South Asian populations. Furthermore, analyses of basal N mtDNA haplogroup (Fernandes, et al. 2012) suggested that the west-east axis can be as ancient as the aftermath of the successful OOA migration, with the older signal located around the then exposed basin of the Arabo-Persian Gulf. Only recently, it has been demonstrated (Fernandes, et al. 2019) that the axis was maintained throughout as testified by distinctive positive selection signals for more recent variants conferring malaria protection (higher in the west) and lactose tolerance (local adaptation in west and European-derived allele in the east).

The scarcity of human ancient remains in AP is extreme, with archaeological sites containing mainly tools and artifacts (Groucutt, et al. 2018; Petraglia, et al. 2012; Petraglia, et al. 2020; Rose 2010; Rose, et al. 2011). Nonetheless, aDNA information from neighbouring regions and from important periods can be paramount in understanding the genetic history of human groups in AP. A main issue is related with the ancestral population group bearing the basal Eurasian lineage, which is necessarily an outgroup in relation to all known deep ancient hunter-gatherer (HG) Eurasian specimens, such as the 45 ka Ust’-Ishim (Fu, et al. 2014) and 39 ka Oase 1 (Fu, et al. 2015). The basal Eurasian bearers diverged from the common European-East Asian pool before 50 ka (Lazaridis, et al. 2014; Yang and Fu 2018), and it is still a matter of discussion where they were geographically located. Higher proportions of the basal Eurasian lineage have been observed in ancient samples from the Caucasus Mountains (Satsurblia and Kotias specimens; 13-9 ka (Jones, et al. 2015)), the Levant and Iran (14-3.5 ka; (Lazaridis, et al. 2016)). To a lesser extent, the lineage is still present in modern populations from western Eurasia and southwest Asia (Lazaridis, et al. 2016). It has been previously speculated that isolated basal Eurasian lineage descendants in the Central Zagros Mountains of Iran originated a farmer population (first goatherds) that spread afterwards into the Eurasian steppe, similarly to the Anatolian-related farmers spread westward into Europe, and Levant farmers spread southward into East Africa. Eventually, by the time of the Bronze Age, the three farmer groups mixed together and with the European hunter-gatherers, reducing the initial genetic differentiation between those groups (Broushaki, et al. 2016; Lazaridis, et al. 2016). Also the ancient Iberomaurusians specimens could be in part descendants of basal Eurasians (van de Loosdrecht, et al. 2018), as they have a shared genetic affinity with early Holocene Near Easterners (best proxy are the Natufians) and one-third of input from sub-Saharan Africans (a mixture between west and east Africans). Thus, so far, southwestern Asia is still the most probable homeland for the hypothetical basal Eurasian population, although a more precise location is still difficult to pinpoint. Another factor to have in mind is the little or no Neanderthal ancestry input in the basal Eurasian lineage (Lazaridis, et al. 2016), supporting a timing for the Neanderthal-modern human interbreeding event after the basal Eurasian lineage divergence and before the separation of the European and Asian pools. The later-on admixture of the basal Eurasian descendants with west Eurasians and southwest Asians possibly explains their observed lower Neanderthal input (Almarri, et al. 2020; Rodriguez-Flores, et al. 2016; Vyas and Mulligan 2019) than in present-day East Asians (no basal Eurasian input). Again, this new information for the Neanderthal-modern human interbreeding event can help to refine the geographic source of the basal Eurasian ancestors. The presence in Israeli archaeological sites of modern humans, such as at Skhul and Qafzeh (Grün and Stringer 1991; Hershkovitz, et al. 2015), and Neanderthals at Tabun (Simpson, et al. 1998) and Kebara Cave (Trinkaus, et al. 1991; Valladas, et al. 1987) dating between 50-120 ka provides evidence that modern humans and Neanderthals overlapped geographically and temporally in the Levant (Hershkovitz, et al. 2015). No such evidence was found in southern parts of southwest Asia, although absence of evidence is not evidence of absence.

In this work, we merged and analysed ancient genomes from key neighbours with the genomic profiles of modern-day Arabian populations in order to test alternative scenarios explaining their origins. Our dataset consisted in 741,000 variants screened in 291 Arabians and 78 Iranians that we previously analysed in detail in terms of population structure and selection events (Fernandes, et al. 2019). Our work differs from others for the good geographical resolution across the entire AP.

## Material and Methods

### Datasets

The “modern dataset” included 291 individuals from AP (100 Saudi Arabia, 61 Yemen, 59 Oman and 71 UAE) and 78 from Iranian populations previously published in Fernandes et al. (Fernandes, et al. 2019). These numbers reflected already only non-related and non-recent migrants from sub-Saharan Africa (inferred in that publication, and reconfirmed here in a Principal Component Analysis). This dataset was merged with 1,998 worldwide relevant available samples (Supplemental Table S1) through PLINK 1.9 (Chang, et al. 2015). Minimum values of 2% were set for missing genotype per marker and missing genotype per sample, leading to a final set of 291,595 SNPs and 2,367 samples. Genotypes from the “modern dataset” were phased with SHAPEITv2.r79044 (Delaneau, et al. 2011) using the 1000 Genomes phased data (Auton, et al. 2015) as a reference panel and using the HapMap b37 genetic map (Frazer, et al. 2007). For algorithms relying on independent markers, SNPs from the modern dataset were pruned for pairwise linkage disequilibrium (LD), by removing any SNP that had an r2>0.2 with another SNP, within a 50-SNPs sliding window with steps of 5 SNPs. After pruning, the “modern dataset pruned” included 116,084 autosomal SNPs and 2,367 individuals. The ancient dataset included 304 samples from several recent studies (downloaded from David Reich Lab website; https://reich.hms.harvard.edu/datasets) as detailed in Supplemental Table S2. The two datasets were merged in Ancient-Modern Dataset (“AMD”) and checked for removal of SNPs with more than 50% of missing genotype (--geno 0.5) and samples with less than 10,000 SNPs, resulting in 293,034 SNPs (or 57,338 SNPs when pruned) and 2,642 samples (275 ancient and 2,367 modern samples).

### Population structure and differentiation

To visualize the genetic similarities of ancient and modern samples, principal component analysis (PCA) was performed in smartpca implemented in EIGENSOFT (Patterson, et al. 2006; Price, et al. 2006) on the AMD. The ancient samples were projected on the modern dataset using the parameters “lsqproject” and “shrinkmode” options turned on.

To analyse the genetic relationships between modern and ancient samples, the ADMIXTURE software (Alexander, et al. 2009) was applied on the “modern dataset pruned” (but limited to the 57,338 SNPs), using maximum likelihood for components (K) from 2 to 20, with the optimal K estimated through cross-validation (CV) of the logistic regression (K=9). The ancient samples were projected on the previously inferred ancestral allele frequencies (K=9).

Population structure was also evaluated on the “modern dataset” based on haplotype information through CHROMOPAINTERv2 (CP) and fineSTRUCTURE v2.07 (Lawson, et al. 2012), in which the modern samples were used both as donors and receivers (all vs all method). The Ne (“recombination scaling constant”) and θ (per site mutation rate) parameters were inferred performing 10 iterations on six randomly selected chromosomes [1,6,10,15,18 and 22] and around 10% of the total samples from each population making a total of 239 samples. The resulting estimates were then averaged taking into account the different recombination pattern of each analysed chromosome. The resulting parameters (Ne=497.390, θ=0.0009938) were then used in the painting runs as input values for -n and -M flag, respectively. The output matrices were merged using the chromocombine tool. The fS was run by using 1,000,000 burn-in and sample iterations, thinned every 10,000 steps and by 100,000 tree comparisons. The maximum concordance tree was built by using both the best posterior and the initial state tree coupled.

RFMix software (Maples, et al. 2013), was used in the modern AP and Iranian samples to infer ancestry for each segment of the genome between a mixture of putative ancestral panels of haplotypes. 1000 Genomes populations were used as parental: Great Britain, representing European ancestry; Yoruba from Nigeria, as the sub-Saharan African ancestry; and Indian Telugu for the South Asian ancestry. Then the median sizes (in cM) of the sub-Saharan African blocks were estimated per AP and Iranian individuals affiliated in clusters A and D.

### Modelling the ancestry composition of AP

QpWave and *qpAdm* tools from the ADMIXTOOLS software package (Patterson, et al. 2012), which relate a set of ‘left’ populations (the population of interest and potential sources of ancestry) to a set of ‘right’ populations (diverse outgroups, Supplemental table S3), were used to test for the number of streams of ancestry from ‘right’ to ‘left’ and estimate mixture proportions. All the possible combinations of N= (2..5) and one set of right/left outgroups were evaluated, as done in (Lazaridis, et al. 2016). For each combination, qpWave was used to evaluate whether the selected outgroups were able to discriminate the combinations of sources, to establish if AP may be explained by the ancient sources and to find the minimum number of ancestry flows between them. Finally, *qpAdm* was used to find the ancestry proportions of the source populations that contributed to the target populations. In both qpWave and *qpAdm* tools, a p-value threshold of 0.01 was set and only feasible mixture proportions were reported.

The Ne and θ values (Ne=497.390, θ=0.0009938) obtained in the CP analysis of the modern dataset were used in an “unlinked” CP analysis with the adaptation of the Non-Negative Least Squares (NNLS) function as in (Leslie, et al. 2015; Montinaro, et al. 2015), by adding ancient samples informative for this geographic region (and having the least number of missing genotypes). CP/NNLS estimates the proportions of the genetic contributions from ancient periods to the modern clusters (Raveane, et al. 2019). For every tested model (per period – Palaeolithic, Neolithic, Bronze Age; overall per country and per cluster – limiting to the four sources identified in the *qpAdm* results), the standard errors were estimated through a weighted jackknife bootstrap, by removing one chromosome at a time, and averaging the values taking into account the total number of markers analysed for iteration (Busing, et al. 1999). In order to evaluate the fitness of the CP/NNLS estimations, the sum of the squared residuals were inferred and the ones with the lowest values were selected as the best estimations for the tested models (Raveane, et al. 2019).

### Impact of the basal Eurasian ancestry

To address the impact of the basal Eurasian lineage in AP (Lazaridis, et al. 2014; Petr, et al. 2019), the *statistic-f4* implemented in the ADMIXTOOLS package (Patterson, et al. 2012) was applied to the modern samples organized per countries and per cluster. The ~45 ka Ust’-Ishim Upper Palaeolithic Siberian (Fu, et al. 2014), who is a proxy for non-basal Eurasian ancestry (Lazaridis, et al. 2016), was considered in the test *f4*(Modern population Test, Han; Ust’-Ishim, Mbuti), as performed in (Fu, et al. 2016). Significant negative values (Z<-3) are evidence for the test populations having ancestry from the basal Eurasian lineage.

Additionally, the *qpAdm* tool was also used to search for traces of the basal Eurasian lineage in modern AP populations. The tested model was in the form (Test, Mota, an early hunter-gatherer-represented by NWRussia - I0061), and one set of right outgroups (Supplemental table S4), following (Lazaridis, et al. 2016). Mota is an eastern African ancient sample (4.5 ka; (Llorente, et al. 2015)) that similarly to the basal Eurasians is basal to other non-Africans, hence indirectly providing an estimate of the mixture proportions of the basal Eurasian lineage. The 7 ka Karelia sample (Haak, et al. 2015) was used as early hunter-gatherer. To relate the basal Eurasian lineage and Neanderthal proportions, the Altai Neanderthal, Denisovan and Mbuti samples were extracted and merged with the modern phased dataset (276,035 SNPs). We then performed a *f4-statistics* analysis in the form *f4*(Test, Mbuti; Altai, Denisova) using the ratio implemented in the ADMIXTOOLS package qpDstat (Patterson, et al. 2012), and the two proportions were correlated as in (Lazaridis, et al. 2016), through a linear regression using the lm function in R.

As the basal Eurasian lineage input in AP could be due to admixture from diverse later already admixed sources, *f4* tests were conducted in the form of *f4*(Test, Han; Ust’-Ishim, X), in which X represented: the Caucasus hunter-gatherer Kotias (9 ka) or Satsurblia (13 ka) (Jones, et al. 2015); ~13 ka Natufian farmer representative of Levant input (Lazaridis, et al. 2016); ~9 ka Zagros farmer (sample ID WC1.SG from (Broushaki, et al. 2016)); ~15 ka North Africa Iberomaurusian (sample TAF011 from (van de Loosdrecht, et al. 2018)).

## Results

### Projection of ancient ancestry on current day Arabian and Iranian diversities

In order to understand how the Arabian and neighbouring populations are related to groups that occupied the region since the transition from the Palaeolithic till the Bronze Age, we performed a PCA projecting ancient samples onto PCs estimated on modern allele-frequency variation (Figure 1). Modern Saudi Arabian and Yemeni samples clustered tightly in the lower left quadrant, overlapping with the three Natufian samples, and were close to the Levant PPNB-PPNC and Levant Bronze Age samples. A part of the modern UAE and Oman samples were more dispersed and mingled with North African-Iberomaurusian and North African-Early Neolithic, in accordance with some sub-Saharan input in their genomes (Supplemental Figure 1 displays a PCA with sub-Saharan and East Asian samples included; Supplemental Figure 2 displays only the modern samples in colours for an easier visualization). While other modern UAE, Oman and the Iranian samples mingled with North African-Late Neolithic, Iran-Late Neolithic and -Neolithic, and Caucasus-Palaeolithic, -Early Bronze Age and -Copper Age. The ancient Anatolian, Balkan, Steppe and European samples were projected distantly from the Arabian and Iranian modern samples.

**Figure 1:**
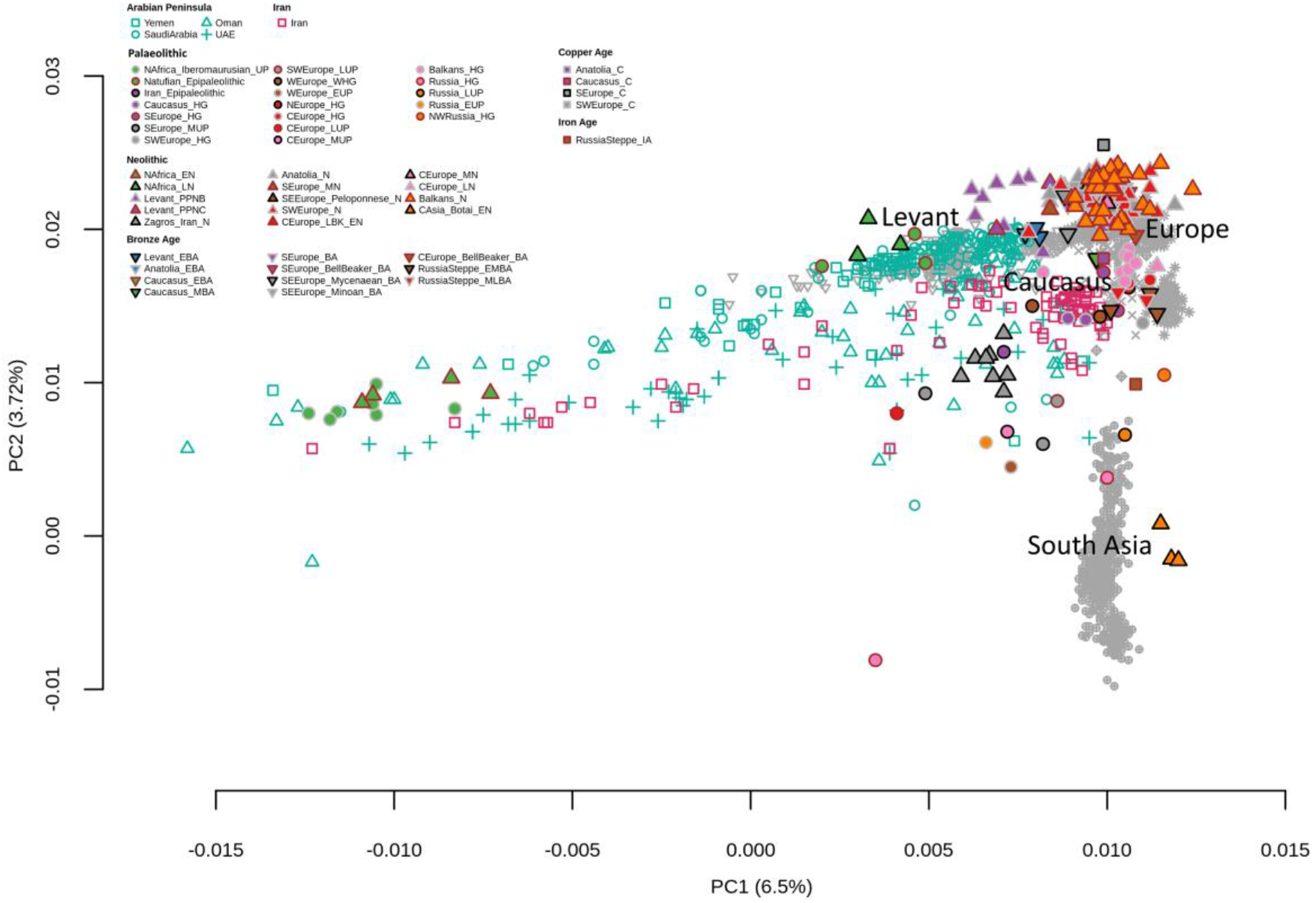
Principal components analysis of present-day samples from the Arabian Peninsula (blue), Iran (red) and neighbouring populations (grey, to avoid visual clutter) with projected ancient samples in different colours and formats according to the legend. Abbreviations are as follows: E, Early; M, Middle; L, Late; HG, hunter-gatherer; N, Neolithic; C, Chalcolithic; BA, Bronze Age; IA, Iron Age; UP, upper Palaeolithic.

The population structure pattern just described for the PCA was largely corroborated by the ADMIXTURE analysis (Figure 2A, B), for K=9. The dark blue component was frequent in modern AP populations (highest values in Saudi Arabia and Yemen) and the Near East, and in the ancient Levant Natufians and Levant Bronze Age samples. In contrast, a substantial proportion of modern samples from East AP and Iran shared a high frequency of the light purple component with Zagros-Neolithic and Anatolia/Caucasus across all periods; this component has been found modal in Levant and Caucasus populations. The orange component, which was very frequent in modern northern European samples, was also dominant in the hunter-gatherer samples from Europe and the Balkans and in the steppe populations. The light green component, the second most common in modern Europeans, especially from the south, and also present in modern Near Easterns, Anatolians and Iranians, was observed in all ancient samples (especially in Anatolian- and Southern European-Neolithic) except in the Zagros-Neolithic ancient sample. In the Zagros-Neolithic ancient sample, as well as in the ancient steppe populations, a considerable amount of the dark green component was detected. This component, typical of modern South Asian populations, was also observed in modern samples from Iran, UAE and Oman. It could be possible that people related to both early farmers from Iran and pastoralists from the Eurasian steppe who are said to have spread eastward into South Asia (Lazaridis, et al. 2016) also had a pronounced impact in eastern AP. The impact of the African ancestry, represented by the pink and red components, was especially high in the ancient South Eastern African samples and has been found with important frequencies in ancient North African individuals (especially in Iberomaurusian samples). The same components were also found in part of modern samples from UAE and Oman, together with a few individuals from Yemen.

**Figure 2:**
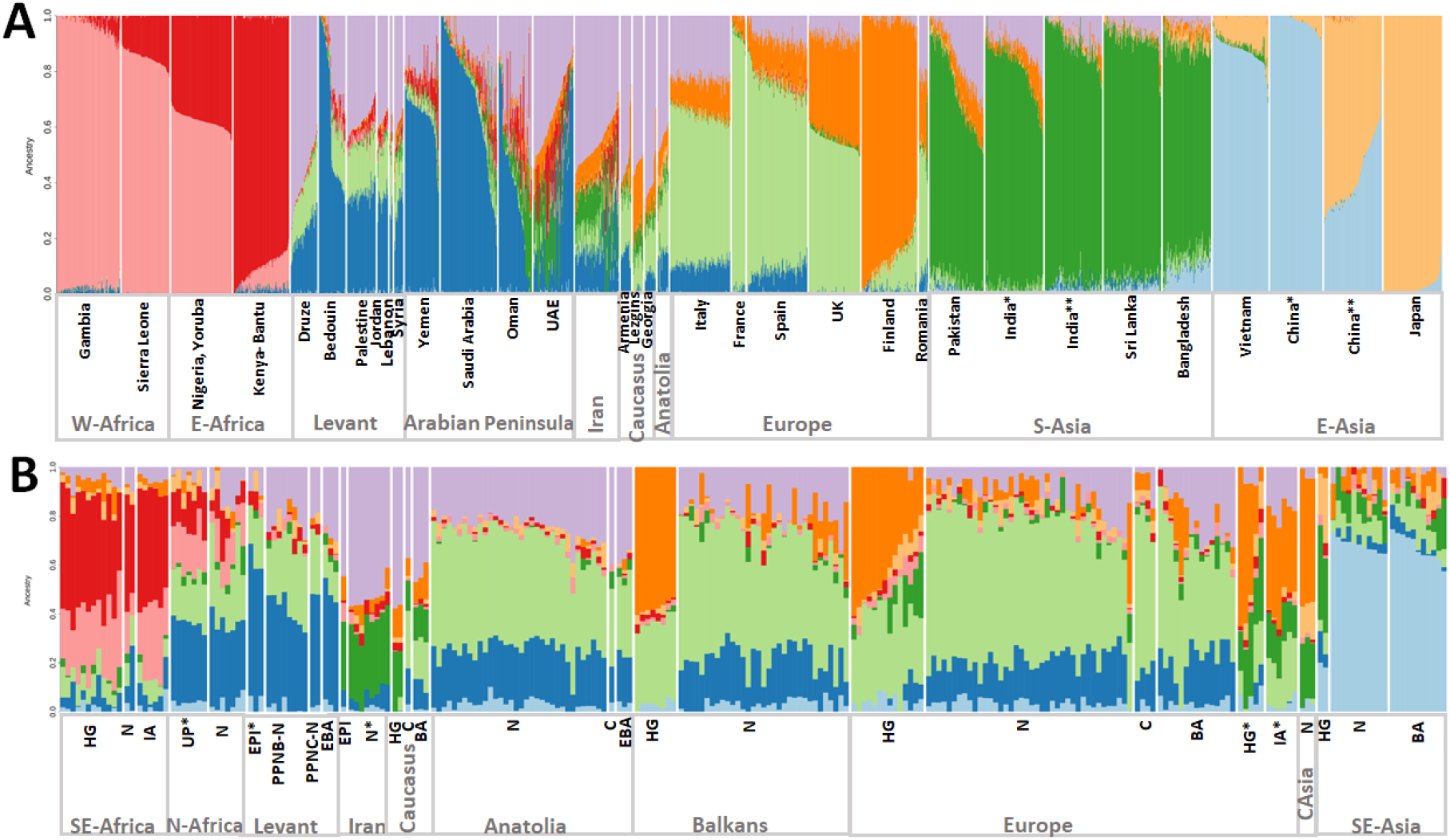
ADMIXTURE model-based clustering analysis of present-day populations (A) and projected ancient samples. India* corresponds to India Gujarati, India** to India Andhra Pradesh, China* to China Xishuangbanna and China** to China Beijing. (B) For the optimal K (K=9) estimated through cross-validation of the logistic regression. UP* corresponds to the Iberomaurusian samples, EPI* to the Natufian samples, N* to the Zagros samples, HG* to samples from Russia and IA* to the steppe samples. Abbreviations are as follows: E, Early; M, Middle; L, Late; HG, hunter-gatherer; N, Neolithic; C, Chalcolithic; BA, Bronze Age; IA, Iron Age; UP, upper Palaeolithic; EPI, Epipaleolithic.

In order to better ascertain the impact of the sub-Saharan African input in the AP genomes (individuals with values > 40% sub-Saharan input had been previously removed from the analysis, to avoid bias by very recent migrants), we checked the population structure through haplotype-based methods (Figure 3A-B; Supplemental Figure 3). A few clusters supported the differentiation between the west-east AP axis: cluster F is typical of west AP, representing 2/3 of Saudi and Yemeni populations, being less than 1/3 frequent in UAE and Oman; cluster D is characteristic of east AP, being also shared with Iran, and practically inexistent in west AP; cluster A, was highly frequent in east AP (nearing 50%), and more restricted in west AP (around 20%). Cluster C, typical from the Levant showed quite limited presence in Saudi, UAE, Oman and Iran; this cluster is related with cluster B only observed in Druze people (an isolated group in Levant). The mean sub-Saharan input values in the modern AP and Iranian samples distributed by these clusters were the following: 22% in A; 5% in C; 6% in D; 0.3% in E; and 3% in F. Definitely, cluster A has still a considerable input from sub-Saharan Africa, but all the other have limited values around 5%. We further tried to ascertain the lengths of the sub-Saharan blocks in clusters A and D (Supplemental Figures 4-5), and there is a clear tendency for the blocks being significantly lower in cluster D, compatible with an older age of admixture than in cluster A.

**Figure 3.**
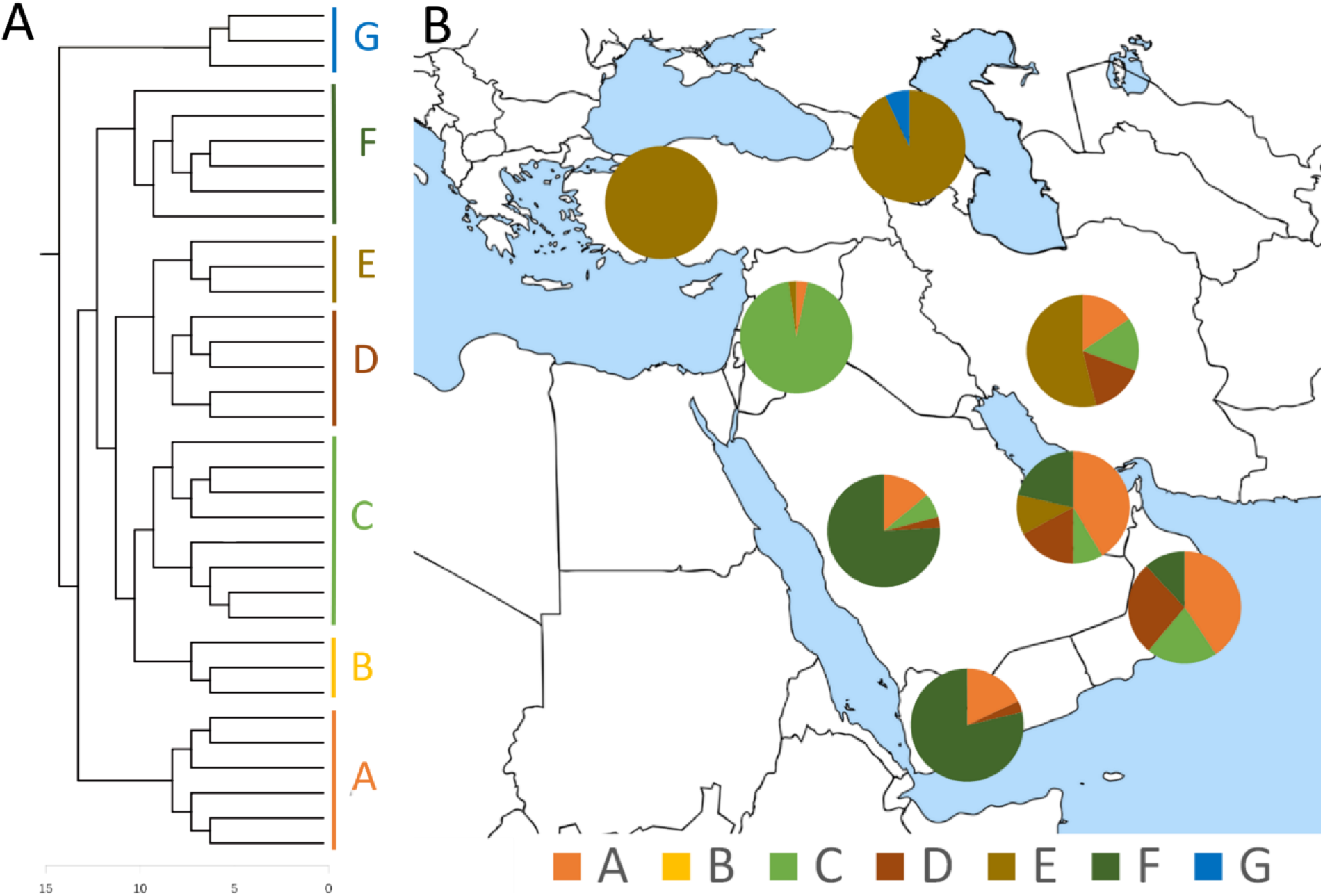
Population substructure in AP and neighbouring regions (see Supplemental Figure 3 for detailed tree). **A.** Finestructure tree with population clusters highlighted. The outgroup G are European populations. Cluster B is made up of only Druze individuals, which were not represented in section B of the figure. **B.** Distribution of the clusters per AP country and in Iran, Levant (summing up Palestinians, Jordanians, Syrians and Lebanese individuals), Caucasus (Armenians, Georgians and Lezgins) and Anatolia (Turkey).

### Modelling the ancestry composition of AP

First we modelled the ancestry composition of AP per country. When using *qpAdm* to model the mixture proportions from ancient source populations the most complex scenario that was feasible for the AP and Iranian populations tested here consisted in four sources (Figure 4A; Supplemental Table S5): Caucasus hunter-gatherer; Iberomaurusian; Natufian; and Zagros farmer. Two thirds of the west AP populations have ancestry shared with the Natufian, typical of the Levant, while east AP and Iran had a predominance of background shared with Caucasus hunter-gatherers and Zagros farmers. The similarity with the Iberomaurusian was higher in east than in west AP, but already residual in Iran. All the other feasible four-source scenarios (Supplemental Table S6) consisted in a predominant Levant signal in the west (replacing either Caucasus or Zagros by Levant PPN or Levant early bronze age), while the east had always a main Caucasus (sometimes Caucasus early bronze age replacing totally the Levant signal) and Zagros farmer similarity.

**Figure 4:**
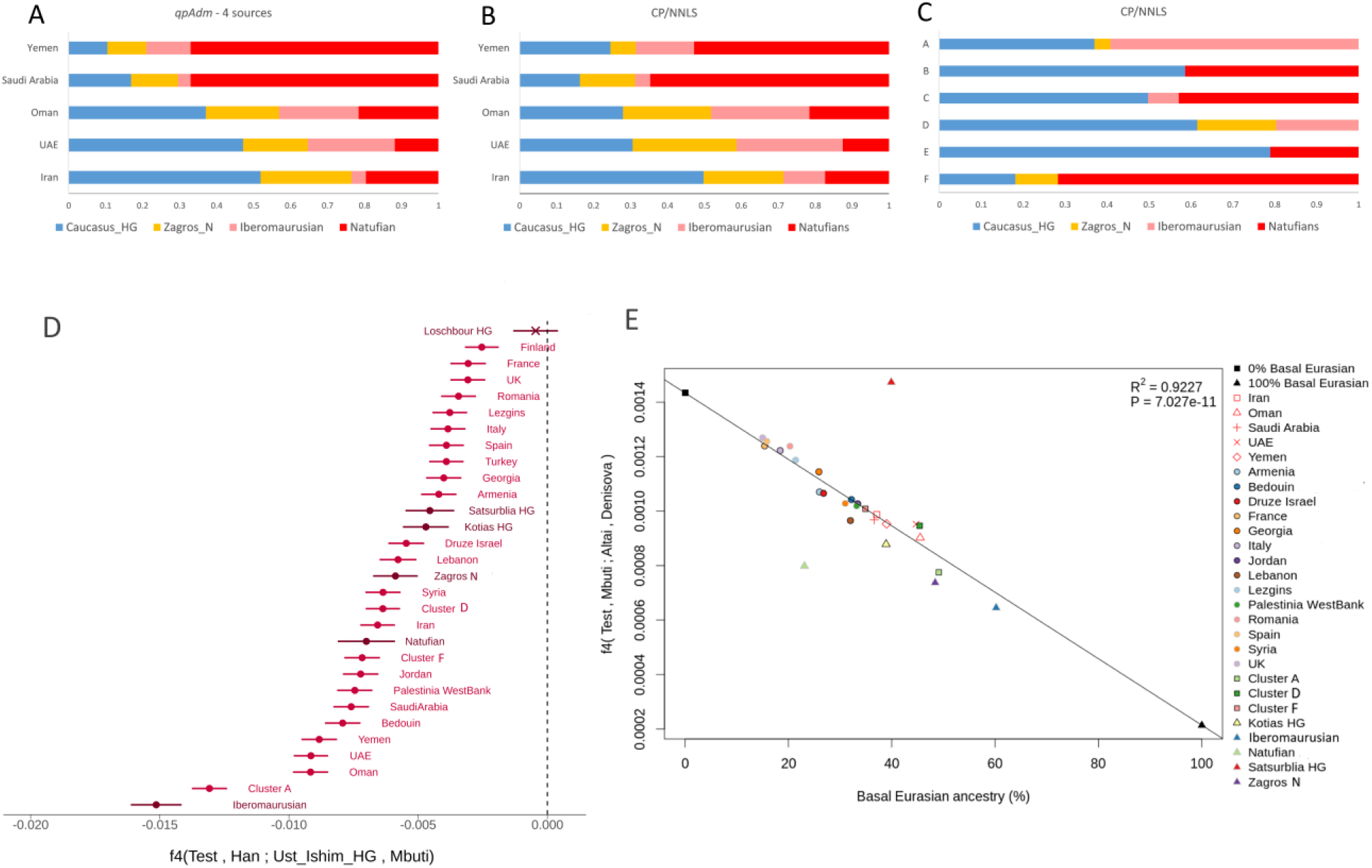
Modelling the ancestry composition of AP. **A.** Mixture proportions on AP and Iran populations inferred by *qpAdm* as a combination of four ancient sources (this combination was the only one feasible in all the five tested populations). **B.** CP/NNLS results for the same set of four ancient sources identified in the *qpAdm* analysis for each country. **C.** CP/NNLS results for the same set of four ancient sources identified in the *qpAdm* analysis for each cluster observed in AP. **D.** Basal Eurasian lineage contribution inferred through the statistic *f4*(Test, Han; Ust’-Ishim, Mbuti), where Ust’-Ishim and Mbuti were used as proxies for non-basal and basal Eurasian lineage, respectively. The bullet identifies significantly negative values (Z-score < -3), while the cross identifies non-significant values, light red indicates modern populations and dark red marks ancient specimens. **E.** Correlation between the basal Eurasian lineage estimates (inferred by *qpAdm*) and the Neanderthal introgression (measured through the *f4*(Test, Mbuti; Altai, Denisovan) test).

The CP/NNLS results per period for the five modern populations studied here (Supplemental Figure 6) again, reflect the similarity within the west AP group and within the east AP-Iran group, with a clear tendency for an older input in the eastern side. As the four source identified in *qpAdm* seem also to be highlighted in the CP/NNLS results per period, we performed an overall CP/NNLS analysis by testing Caucasus hunter-gatherer, Iberomaurusian, Natufian; and Zagros farmer as sources (Figure 4B; Supplemental Table S7). As shown in the figure, relative proportions of each of these sources are quite parallel between *qpAdm* and CP/NNLS inferences (linear correlation; r^2^=0.8632), with a slight exception for Yemen, with a lower Zagros input and a higher Caucasus hunter-gatherer signal.

Given the high frequency in east AP of cluster A that has a significant sub-Saharan African input, we also calculated the CP/NNLS with the four sources (Caucasus hunter-gatherer; Iberomaurusian; Natufian; and Zagros farmer) per cluster (Figure 4C; Supplemental Table S8). Only three clusters had input from Iberomaurusian: this input was overwhelming, and probably biased by a recent influence, in cluster A; it was very restricted in the east AP/Iranian-cluster D and more so in the Levant-cluster C. Curiously, the west AP-cluster F had significantly more influence from Natufian than the Levant –C, Druze-cluster B, and Anatolian/Caucasus-cluster E. Zagros influence was restricted to the east AP/Iranian-cluster D, west AP-cluster F and cluster A.

### Basal Eurasian lineage

The *f4* test with the Ust’-Ishim as proxy for the non-basal Eurasian lineage (Figure 4D) showed significant negative results in most of the tested modern populations from southwest Asia, compatible with the presence of the basal Eurasian lineage in these populations. The significant results were more extreme in east AP followed by west AP, the Near Eastern populations and Iran, Caucasus, Anatolia, and finally Europe. The ancient Iberomaurusian sample included in the analysis was the one with the highest input of the basal Eurasian lineage, even higher than the ones observed in modern east AP populations. Not surprisingly, cluster A bearing still a mean 20% sub-Saharan African admixture, was in a close position to the Iberomaurusian sample, with clusters D (east AP-Iran) and F (west AP) occupying a position similar to the other Levant and Iranian samples. The other ancient samples displayed similar *f4* statistic values to local modern-day populations. The European Louschbour hunter-gatherer specimen, as expected, showed no input from the basal Eurasian lineage.

In accordance with previous reports for ancient samples (Lazaridis, et al. 2016; Vyas and Mulligan 2019), the proportion of the basal Eurasian lineage in the modern samples was negatively correlated with the Neanderthal input (r2=0.92; p=7.03×10^−11^; Figure 4E; (Supplemental Tables S9 and S10). Concordantly with the above results, the modern samples with the highest amount of the basal Eurasian lineage were the east AP populations (~45%), followed by the west AP and Iran (~38%), Levant (~32%; except Druze which displayed a lower 28%) and Caucasus (20-25%), while Europeans had the lowest values (<20%). In this analysis, the basal Eurasian lineage proportions between clusters A and D were quite similar (49% and 45%, respectively), indicating that the bias introduced by a probable recent sub-Saharan input in AP is very low. These proportion of the basal Eurasian lineage in the clusters more abundant in east AP were higher than the value observed for the dominant cluster in west AP (cluster F; 35%). The ancient specimens follow the regression line (the two outliers have higher missing genotypes), with the Iberomaurusian specimen displaying the highest basal Eurasian and the lowest Neanderthal proportions, as expected from the previous results. The *qpAdm*-based inference for the basal Eurasian input indicates comparable amounts in modern AP and ancient samples from Caucasus and Zagros, with a lower input in the Natufian.

When testing *f4* by replacing an indirect modern day basal Eurasian source with an admixed ancient sources (Supplemental Figure 7), we observed a preponderant signature of the Natufian in west AP, that was nevertheless low impacting in east AP (in fact, it had a higher influence in southern European populations than here). The Iberomaurusian had a main signal in the Levant and west AP, concordant with a high Iberomaurusian-Natufian similarity (van de Loosdrecht, et al. 2018), followed by east AP (probably the basal Eurasian similarity), and then Iberia and Italy (which is an interesting result for the discussion of North Africa-South Europe connections). The Caucasus and Zagros specimens, themselves displaying several signals of admixture involving the periods under analysis (Jones, et al. 2015), had a less clear pattern, so that founder and receptor becomes already too complex to disentangle.

## Conclusions

The molecular dating of past admixture events based only on present-day human genomic data is limited to a 5 ka time span due to the constraints caused by recombination (Hellenthal, et al. 2014). Fortunately, as we have shown in this work, the availability of informative aDNA samples from neighbouring key geographic regions allows to directly test alternative scenarios for past founder contributions to the genomic pools of extant populations.

It is now evident the continuum along time linking east AP with Iran and Caucasus, as reflected by the outcomes of the various analyses performed here. We further extend the conclusions reported by others (Gallego-Llorente, et al. 2016; Lazaridis, et al. 2016) of two separated hunter– gatherer to farmer transitions, one in the southern Levant and the other in southern Caucasus–Iran highlands. Our results reveal that the eastern continuum was probably older, given the stronger signal of the basal Eurasian lineage and Iberomaurusians in that side of the AP. Definitely, the Iberomaurusian specimens are the ones bearing a very high level of basal Eurasian lineage, which could have been contributed by the sub-Saharan African input added by the southwest Asian founder group, which moved to North Africa 20 ka bearing a high basal Eurasian component. It is interesting that the current-day southwest Asian populations that show more genetic sharing with Iberomaurusians are the east AP populations. We tried to disentangle between an ancient link and a bias due to recent sub-Saharan input in this signature by performing the haplotype-based clustering, and confirmed a still high proportion of the basal Eurasian lineage/Iberomaurusian founding effect in clusters D (east AP and Iran) and F (west AP). Probably, the Iberomaurusian specimens are currently the best proxy for the basal Eurasian population group.

The highest levels of basal Eurasian lineage found in east AP reinforce previous genetic and archaeological evidence of early human settlement in the exposed basin of the Arabo-Persian Gulf (Al-Abri, et al. 2012; Alshamali, et al. 2009; Cerný, et al. 2011; Fernandes, et al. 2012; Fernandes, et al. 2015; Rodriguez-Flores, et al. 2016; Vyas, et al. 2017). These results, by dislocating the geographic origin of the basal Eurasian population group from the Near East to the eastern part of the Arabian Peninsula, reconcile better with the Neanderthal-modern human interbreeding event having occurred probably in the Levant. Lazaridis et al. (Lazaridis, et al. 2016), who observed that the aDNA specimens with higher basal Eurasian lineage were from the Levant, suggested that either basal Eurasians thoroughly admixed into the Near East necessarily before the date of the analysed Natufians but after the Neanderthal admixture, or the ancestors of basal Eurasians lived in the Near East, but did not participate in the Neanderthal admixture. Our results seem to support the first scenario, although they do not exclude other hypotheses. Thus, the exposed basin of the Arabo-Persian Gulf is a possible homeland of basal Eurasians, with an easy corridor linking the current Hormuz Strait to the Zagros and Caucasus steppes (the east continuum). Recent archaeological evidence (Heydari-Guran and Ghasidian 2020) is being collected in the Zagros region, proving that this region was passable in the Middle and Upper Palaeolithic, and included intermountain plains connected to each other by valleys associated with permanent water and raw material sources. If the basal Eurasians were located in the Arabo-Persian Gulf, another non-African group would be mingling at a northern location, around Levant, with the autochthonous Neanderthals, and were the ancestors of the current European and Asians pools. The basal Eurasians and the Neanderthal admixed group were genetically close, so they most likely descended from the same African migrant group that did split somewhere. Did the split occur in southwest Asia after the OOA migration (through either the northern or the southern route (Lahr and Foley 1994))? Or, alternatively, are our data suggesting that the group split earlier in Africa? In the latter scenario, the subset that gave rise to the basal Eurasian branch probably followed the southern route taking refugium in the exposed basin of the Arabo-Persian Gulf, while the direct ancestors of Europeans and Asians followed the northern route, mixed with Neanderthals, and hence moved forward further splitting towards Europe and Asia. Current evidence does not allow us to disentangle between these scenarios, and highlights the urgency of finding and analysing ancient human specimens in east AP/Zagros region. These specimens do not need to be as old as 50 ka, as 20-10 ka specimens from east AP/Zagros should expectably bear a higher basal Eurasian ancestry and represent excellent proxies for the demographic, admixture and dispersal events on the awake of OOA and around the split of the main population groups existing nowadays.

## Acknowledgments

This work was financed by FEDER-Fundo Europeu de Desenvolvimento Regional funds through COMPETE 2020-Operacional Programme for Competitiveness and Internationalisation (POCI), Portugal 2020, by Portuguese funds through FCT-Fundação para a Ciência e a Tecnologia, Ministério da Ciência, Tecnologia e Inovação in the framework of the project “Biomedical anthropological study in Arabian Peninsula based on high throughput genomics” (POCI-01-0145-FEDER-016609), the Italian Ministry of Education, University and Research project Dipartimenti di Eccellenza Program (2018–2022) - Department of Biology and Biotechnology “L. Spallanzani,” University of Pavia (to A.T.), the Italian Association for Cancer Research (AIRC) and the Italian Ministry of Health (A.R.). VF has a postdoc grant through FCT (SFRH/BPD/114927/2016). i3S is financed by FEDER-COMPETE 2020, Portugal 2020 and by Portuguese funds through FCT in the framework of the project ‘Institute for Research and Innovation in Health Sciences’ (POCI-01-0145-FEDER-007274). Authors would like to thank Dr. Francesco Bertolini for facilitating the research of A.R. in the last stage of the manuscript preparation.

## Notes

### Competing Interest Statement

The authors have declared no competing interest.

