## Supplemental Material for "Projecting ancient ancestry in modern-day Arabians and Iranians: a key role of the past exposed Arabo-Persian Gulf on human migrations"

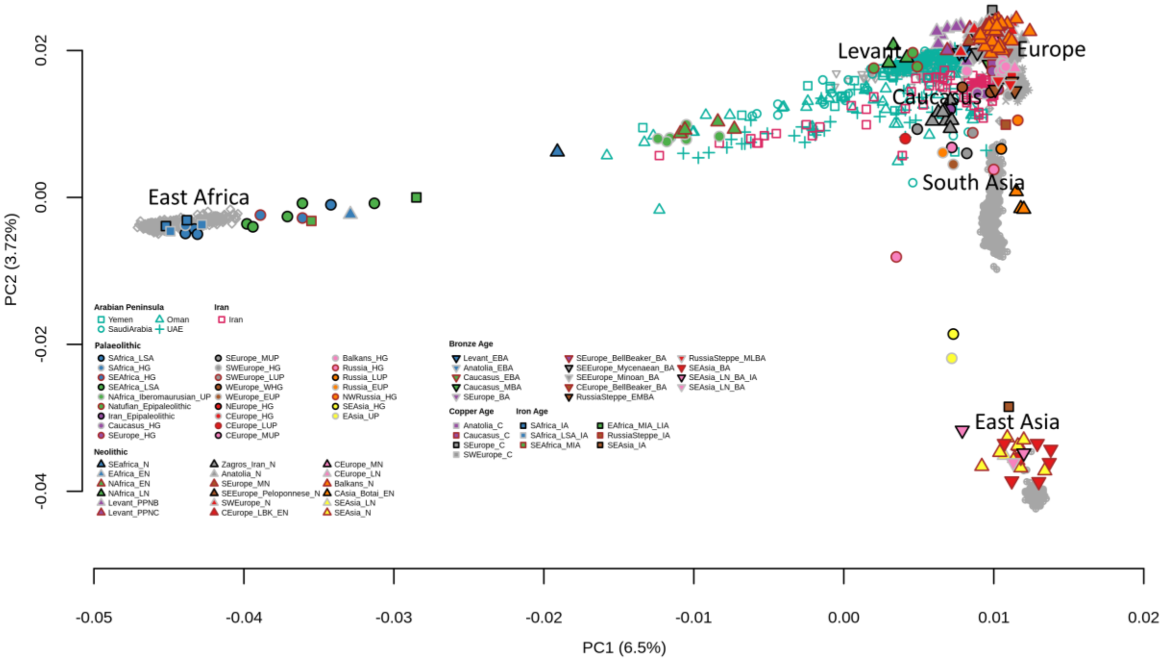

**Supplemental Figure 1**: Principal components analysis of present-day samples from the Arabian Peninsula (blue), Iran (red) and neighbouring populations (grey, to avoid visual clutter) with projected ancient samples in different colours and formats according to the legend. Abbreviations are as follows: E, Early; M, Middle; L, Late; HG, hunter-gatherer; N, Neolithic; C, Chalcolithic; BA, Bronze Age; IA, Iron Age; UP, upper Palaeolithic; EPI, Epipaleolithic.

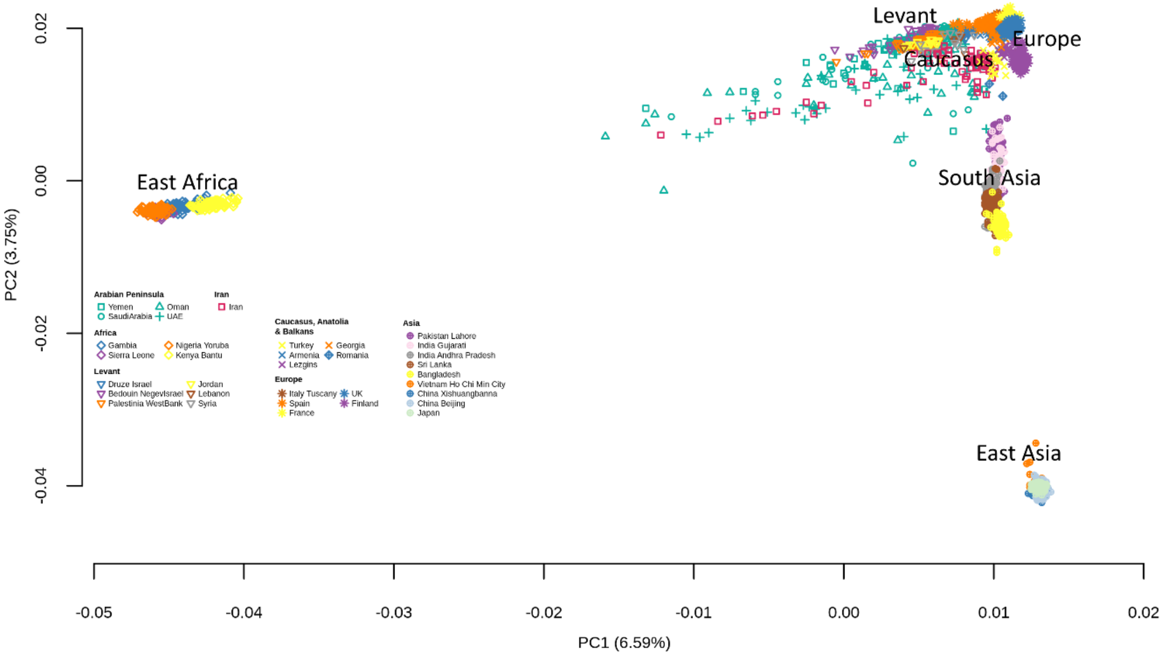

**Supplemental Figure 2**: Principal components analysis of present-day individuals.

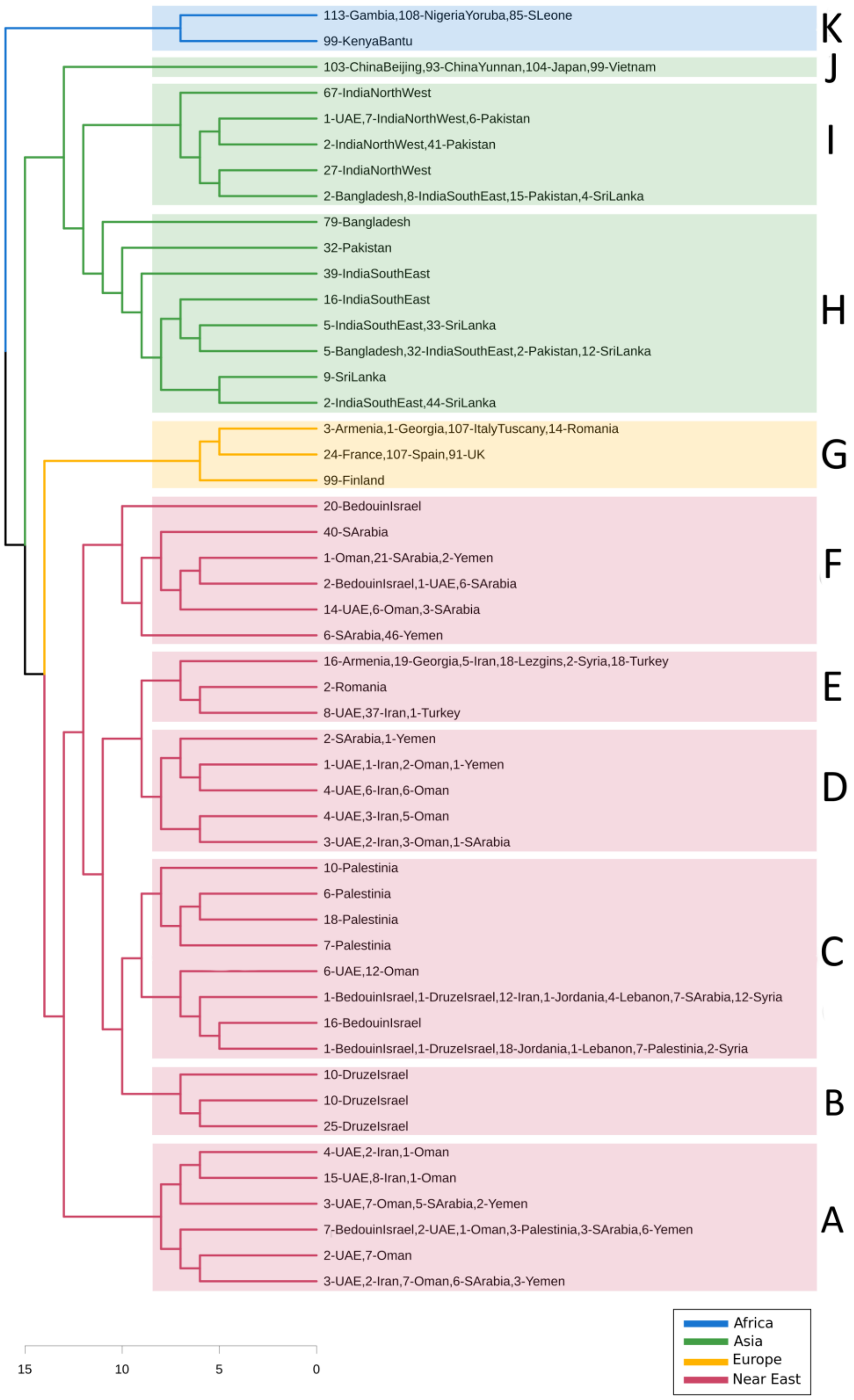

**Supplemental Figure 3**: Hierarchical clustering tree from which were clusters based on sample geographical locations were inferred.

### Iran

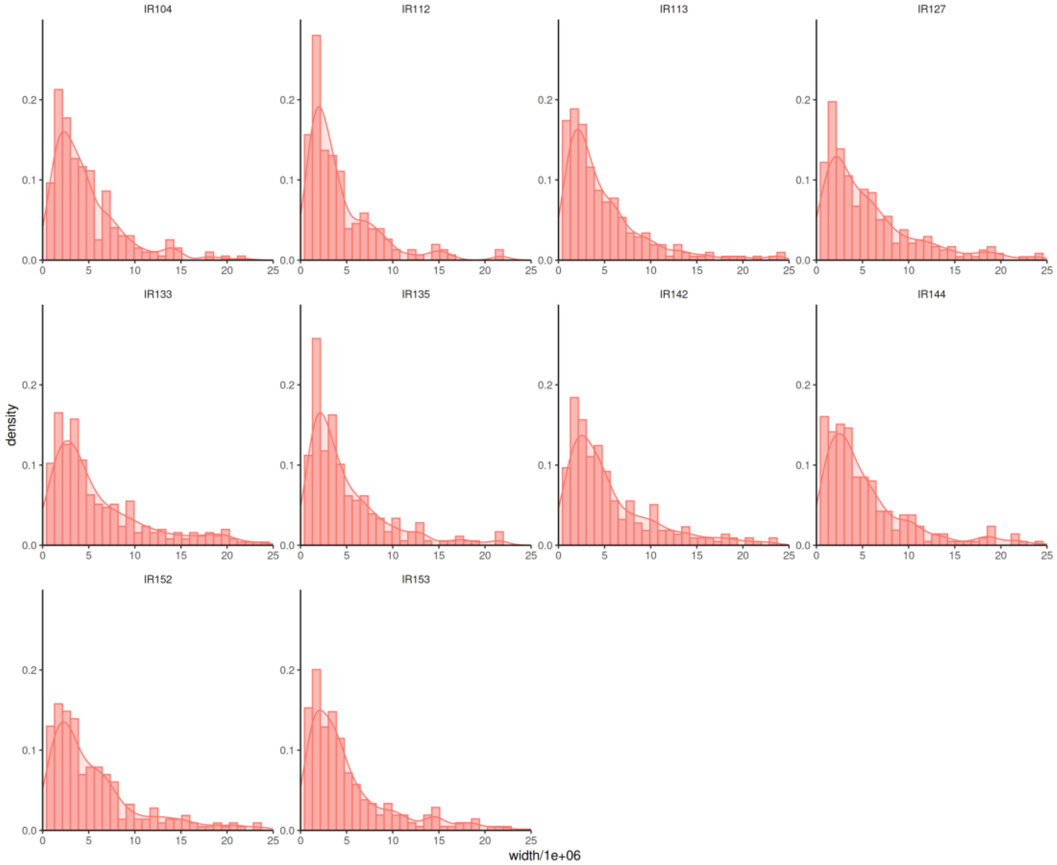
IR104
IR112
IR113
IR127
IR133
IR135
IR142
IR144
IR152
IR153
density
width/1e+06

### Oman

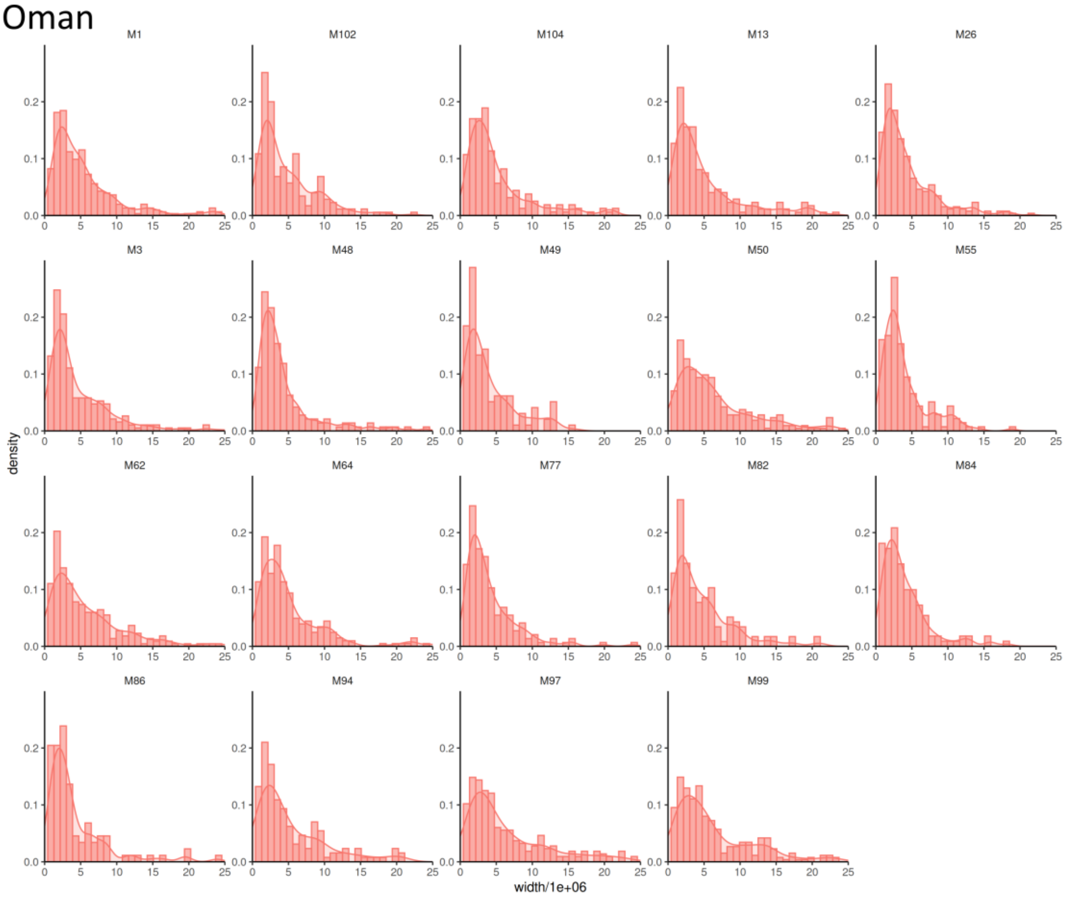
M1
M102
M104
M13
M26
M3
M48
M49
M50
M55
M62
M64
M77
M82
M84
M86
M94
M97
M99
density
width/1e+06

### UAE

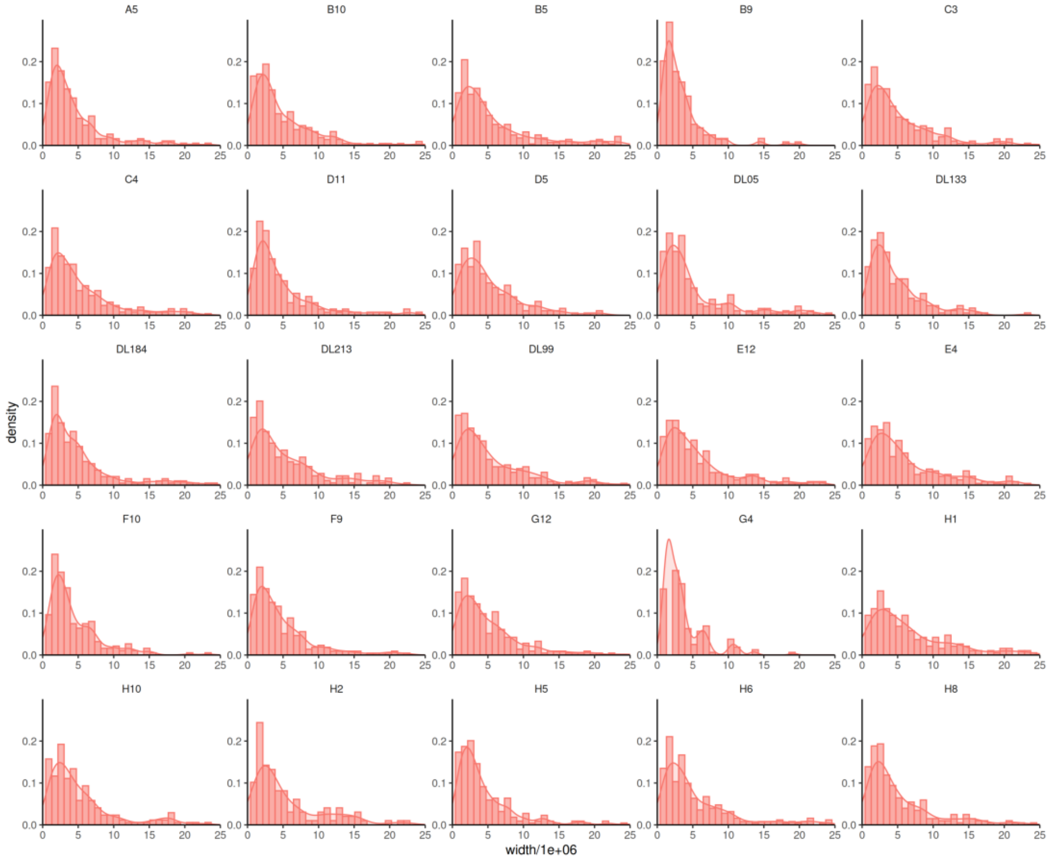

### Saudi Arabia

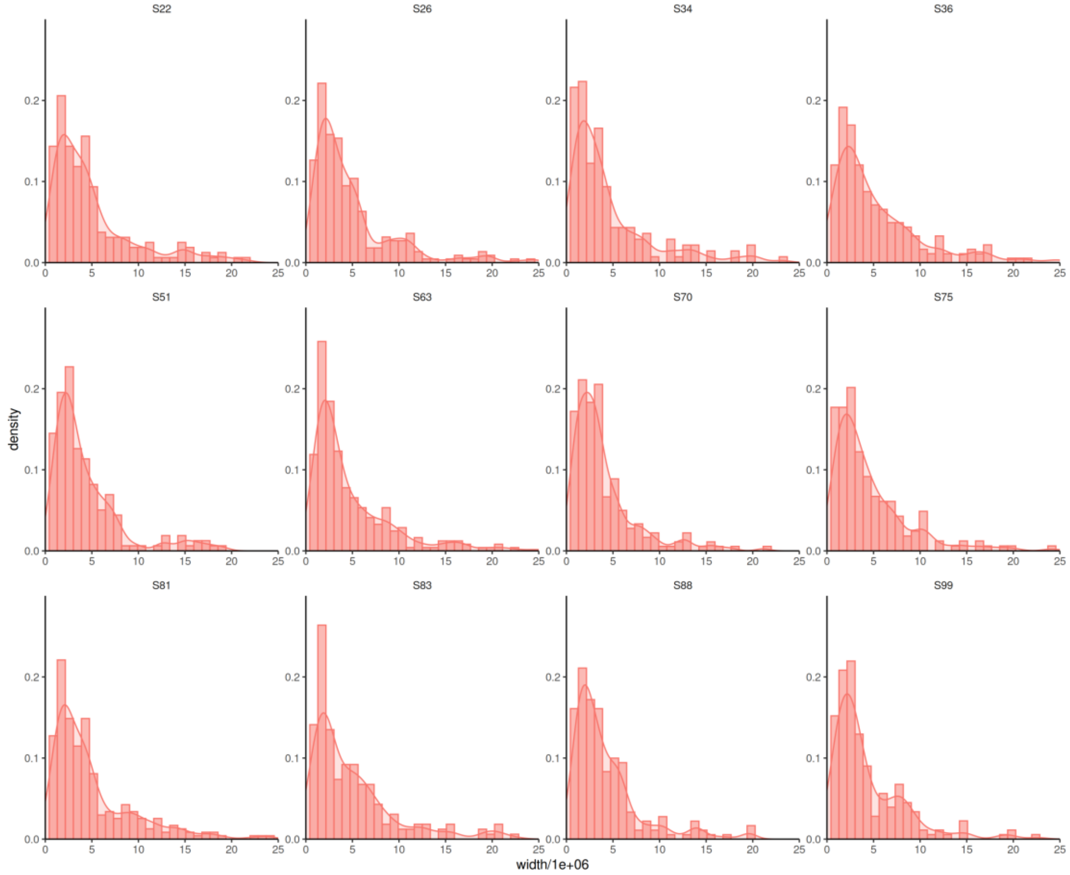

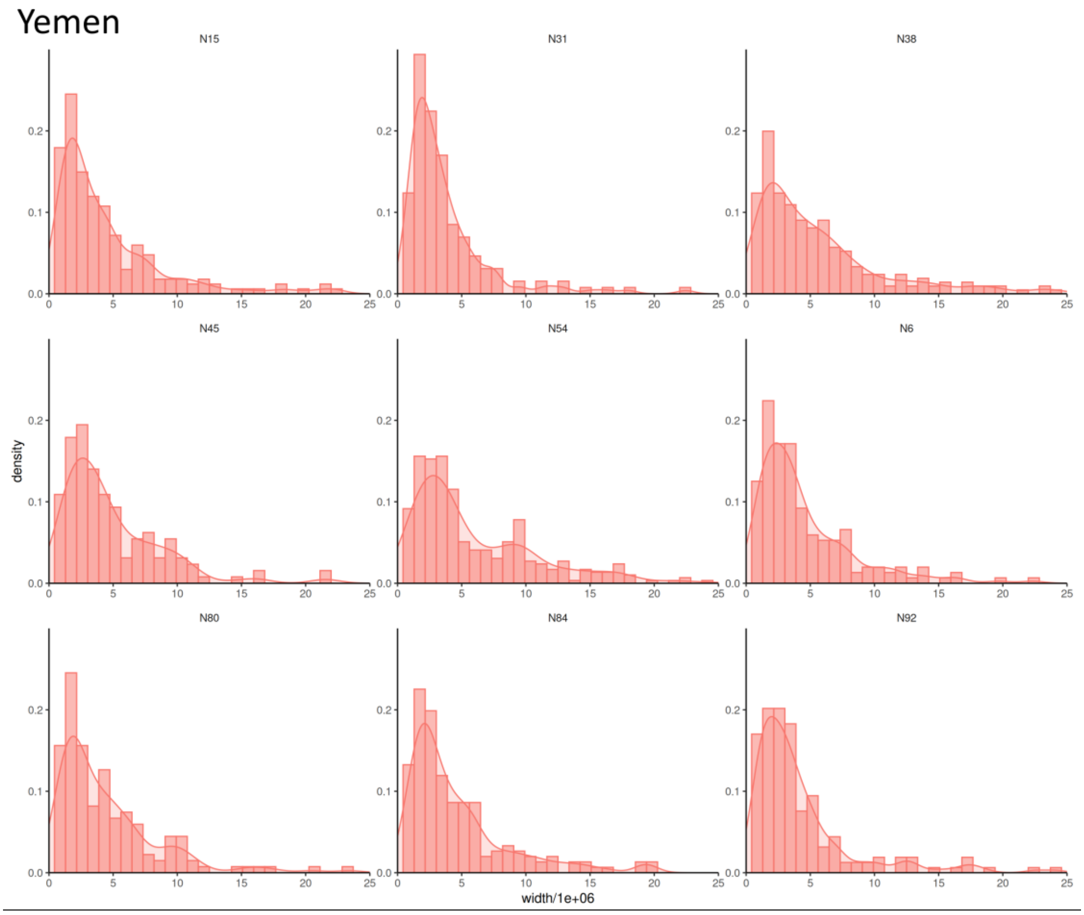

**Supplemental Figure 4**: Median sizes (in cM) of the sub-Saharan African blocks estimated per AP and Iranian individuals affiliated in cluster A.

### Iran

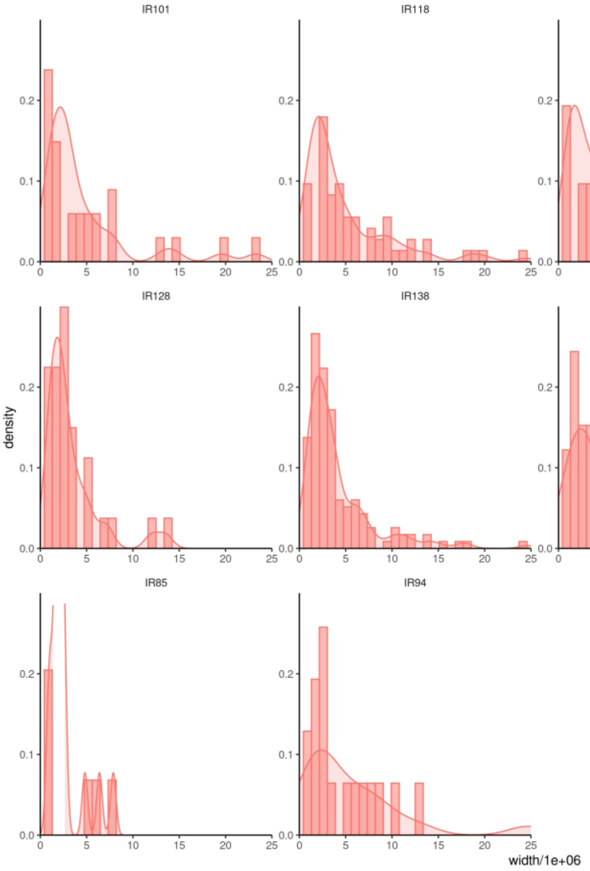
IR101
IR118
IR120
IR121
IR128
IR138
IR149
IR83
IR85
IR94
density
width/1e+06
0.0
0.1
0.2
0
5
10
15
20
25

### Oman

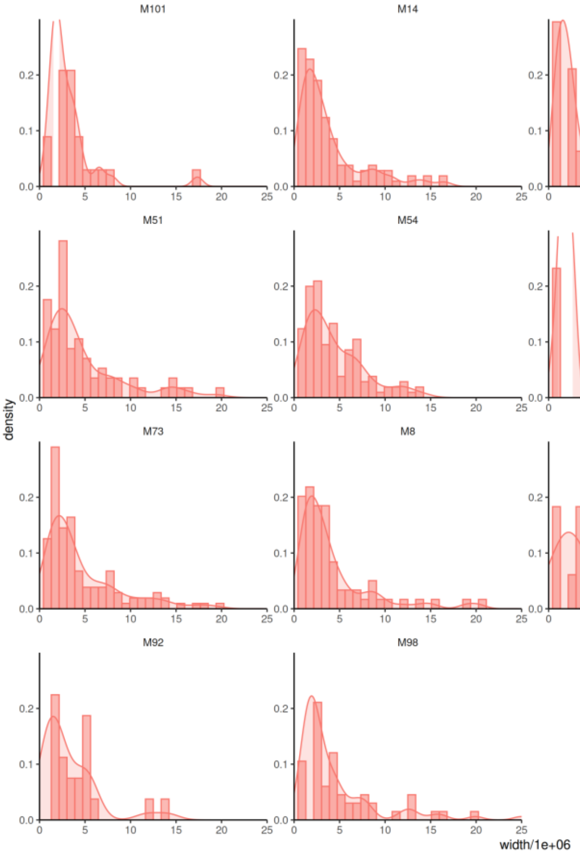
M101
M14
M35
M5
M51
M54
M61
M63
M73
M8
M88
M90
M92
M98
density
width/1e+06
0.0
0.1
0.2
0
5
10
15
20
25

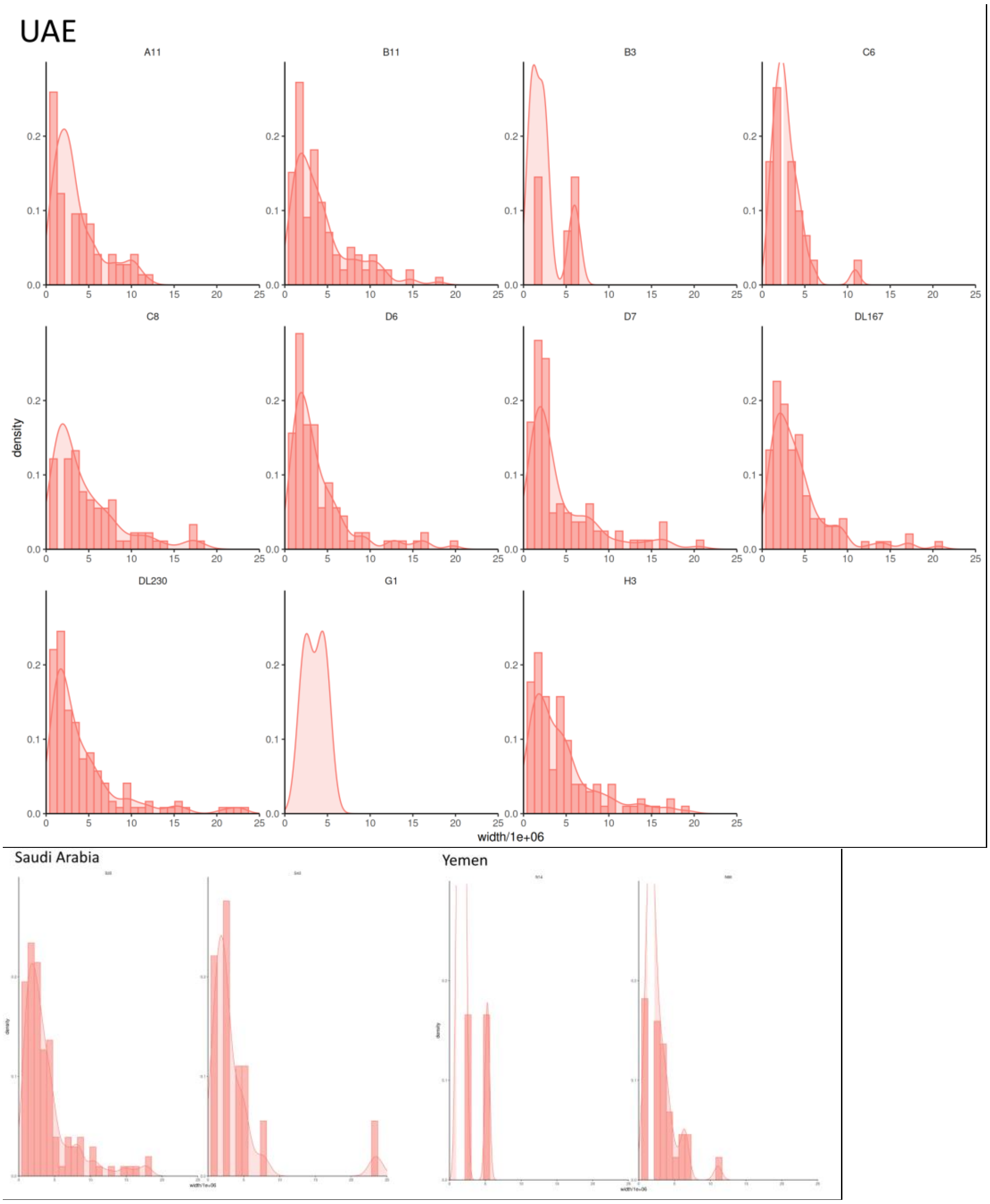

**Supplemental Figure 5**: Median sizes (in cM) of the sub-Saharan African blocks estimated per AP and Iranian individuals affiliated in cluster D.

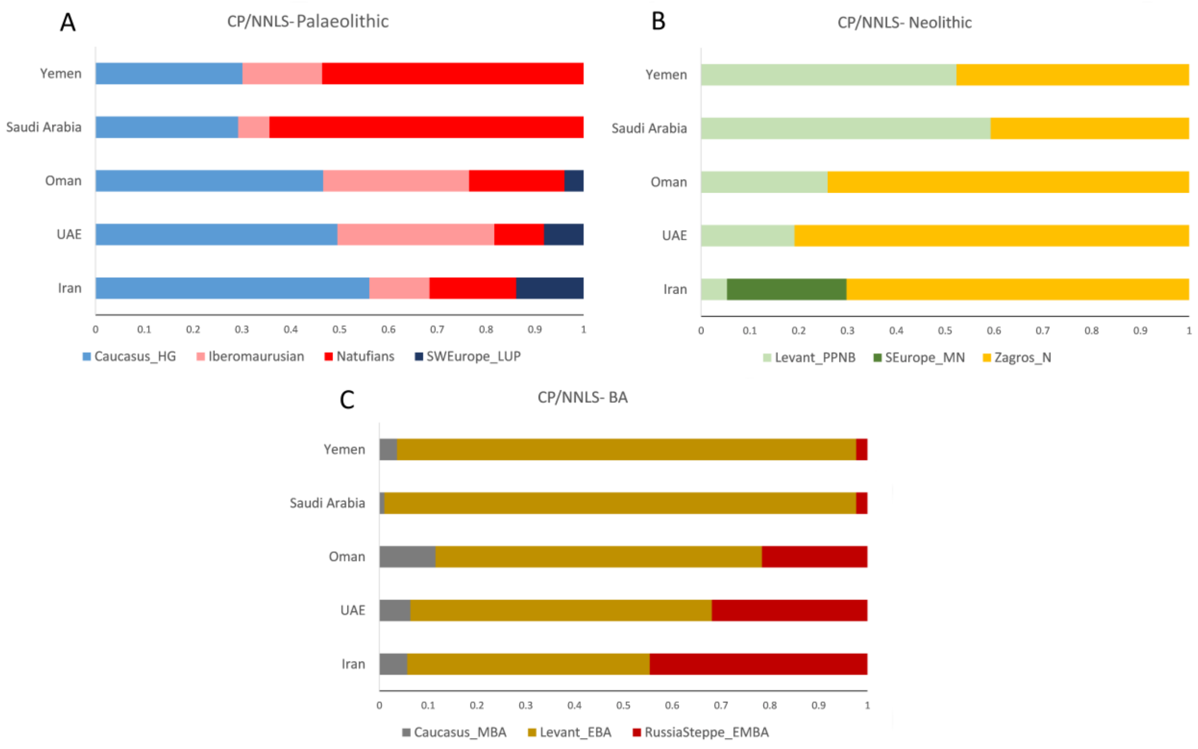

**Supplemental Figure 6**: Population-based ancestry estimates for all the modern populations inferred by CHROMOPAINTER via an NNLS-based analysis. A, Palaeolithic period; B, Neolithic period; C, Bronze age period. Abbreviations are as follows: E, Early; M, Middle; L, Late; HG, hunter-gatherer; N, Neolithic; BA, Bronze Age; UP, upper Palaeolithic.

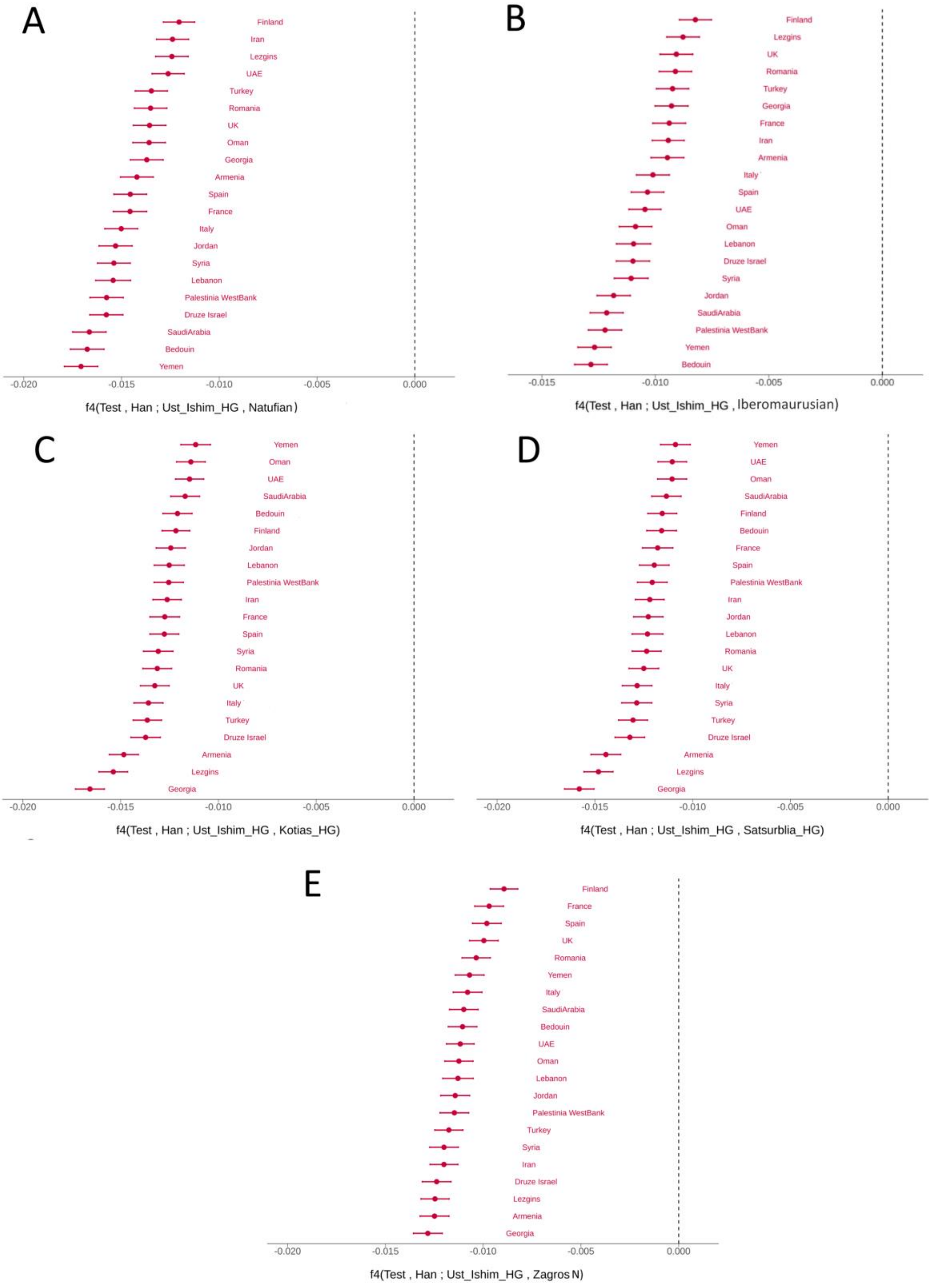

**Supplemental Figure 7**: Alternative testing of the statistic *f4*(Test, Han; Ust'-Ishim, X), where Ust'-Ishim and X were used as proxies for non-basal and basal Eurasian lineage, respectively. The X was a rather recent ancient specimen of admixed basal Eurasian ancestry:

Levant pre-Neolithic Natufian (A); North Africa Iberomaurusian (B) Caucasus Kotias hunter-gatherer (C); Caucasus Satsurblia hunter-gatherer (D); and Iranian Zagros (E) farmer. The bullet identifies significantly negative values (Z-score < -3). All the tested samples were modern populations.

**Table S1:** Modern samples used in this study.

| Country | Region | N. of Samples | Latitude | Longitude | Reference |
|---|---|---|---|---|---|
| Armenia | Caucasus | 19 | 40.11 | 44.31 | Behar et al. (2010) doi: 10.1038/nature09103 and Yunusbayev et al. (2012) doi:10.1093/molbev/msr221 |
| Bangladesh | South Asia | 87 | 23.42 | 90.21 | 1000 Genomes Project Consortium (2015) doi:10.1038/nature15393 |
| Bedouin | Levant | 48 | 30.5 | 34.91 | Patterson et al. (2012) doi: 10.1534/genetics.112.145037 and Lazaridis et al. (2014) doi: 10.1038/nature13673 |
| China Beijing | East Asia | 103 | 39.55 | 116.23 | 1000 Genomes Project Consortium (2015) doi: 10.1038/nature15393 |
| China Xishuangbanna | East Asia | 99 | 22 | 100.48 | 1000 Genomes Project Consortium (2015) doi: 10.1038/nature15393 |
| Druze Israel | Levant | 47 | 32.57 | 35.1 | Patterson et al. (2012) doi: 10.1534/genetics.112.145037; Haber et al. (2016) doi: 10.1038/ejhg.2015.206 and Lazaridis et al. (2014) doi: 10.1038/nature13673 |
| Finland | North Europe | 100 | 60.1 | 24.56 | 1000 Genomes Project Consortium (2015) doi: 10.1038/nature15393 |
| France | West Europe | 24 | 48.51 | 2.21 | Patterson et al. (2012) doi: 10.1534/genetics.112.145037 and Lazaridis et al. (2014) doi: 10.1038/nature13673 |
| Gambia | West Africa | 113 | 13.28 | -16.36 | 1000 Genomes Project Consortium (2015) doi: 10.1038/nature15393 |
| Georgia | Caucasus | 20 | 41.43 | 44.47 | Yunusbayev et al. (2012) doi:10.1093/molbev/msr221 and Behar et al. (2010) doi: 10.1038/nature09103 |
| India Andhra Pradesh | South Asia | 103 | 16.5 | 80.64 | 1000 Genomes Project Consortium (2015) doi: 10.1038/nature15393 |
| India Gujarati | South Asia | 106 | 23.13 | 72.41 | 1000 Genomes Project Consortium (2015) doi: 10.1038/nature15393 |
| Iran | Iran | 100 | 35.41 | 51.25 | Fernandes et al. (2019) doi:10.1093/molbev/msz005; Behar et al. (2013) doi:10.3378/027.085.0604 and Yunusbayev et al. (2015) doi: 10.1371/journal.pgen.1005068 |
| Italy | South Europe | 108 | 43.46 | 11.15 | 1000 Genomes Project Consortium (2015) doi: 10.1038/nature15393 |
| Japan | East Asia | 105 | 35.41 | 139.46 | 1000 Genomes Project Consortium (2015) doi: 10.1038/nature15393 |
| Jordan | Levant | 20 | 31.57 | 35.56 | Patterson et al. (2012) doi: 10.1534/genetics.112.145037 and Lazaridis et al. (2014) doi: 10.1038/nature13673 |
| Kenya | East Africa | 111 | 0.37 | 34.46 | 1000 genomes Project Consortium (2015) doi: 10.1038/nature15393 |
| Lebanon | Levant | 7 | 33.54 | 35.32 | Behar et al. (2010) doi: 10.1038/nature09103 |
| Lezgins | Caucasus | 18 | 43.06 | 46.53 | Behar et al. (2010) doi: 10.1038/nature09103 and Behar et al. (2013) doi:10.3378/027.085.0604 |
| Nigeria Yoruba | West Africa | 110 | 7.23 | 3.55 | 1000 Genomes Project Consortium (2015) doi: 10.1038/nature15393 |
| Oman | Arabian Peninsula | 100 | 23.36 | 58.33 | Fernandes et al (2019) doi:10.1093/molbev/msz005 |
| Pakistan Lahore | South Asia | 97 | 31.32 | 74.2 | 1000 Genomes Project Consortium (2015) doi: 10.1038/nature15393 |
| Palestinia WestBank | Levant | 51 | 31.47 | 35.13 | Patterson et al. (2012) doi: 10.1534/genetics.112.145037 and Lazaridis et al. (2014) doi: 10.1038/nature13673 |
| Romania | Balkans | 16 | 44.25 | 26.06 | Behar et al. (2010) doi: 10.1038/nature09103 |
| Saudi Arabia | Arabian | 120 | 24.39 | 46.46 | Fernandes et al. (2019) doi:10.1093/molbev/msz005 |

| | | | | | |
|---|---|---|---|---|---|
| | Peninsula | | | | |
| Sierra Leone | West Africa | 86 | 8.29 | -13.14 | 1000 Genomes Project Consortium (2015) doi: 10.1038/nature15393 |
| Spain | Southwest Europe | 107 | 40.26 | -3.42 | 1000 Genomes Project Consortium (2015) doi: 10.1038/nature15393 |
| Sri Lanka | South Asia | 105 | 6.56 | 79.52 | 1000 Genomes Project Consortium (2015) doi: 10.1038/nature15393 |
| Syria | Levant | 16 | 33.3 | 36.18 | Behar et al. (2010) doi: 10.1038/nature09103 |
| Turkey | Anatolia | 19 | 39.56 | 32.52 | Behar et al. (2010) doi: 10.1038/nature09103 |
| UAE | Arabian Peninsula | 120 | 25.15 | 55.17 | Fernandes et al. (2019) doi:10.1093/molbev/msz005 |
| UK | Northwest Europe | 93 | 51.3 | -0.7 | 1000 Genomes Project Consortium (2015) doi: 10.1038/nature15393 |
| Vietnam | South Asia | 101 | 10.48 | 106.39 | 1000 Genomes Project Consortium (2015) doi: 10.1038/nature15393 |
| Yemen | Arabian Peninsula | 61 | 15.2 | 44.12 | Fernandes et al. (2019) doi:10.1093/molbev/msz005 |

**Table S2:** Ancient samples used in this study.

| SampleID | Group ID | Country | Time Period | Average of 95.4% date range in calBP (defined as 1950 CE) | Sex | Latitude | Longitude | Publication | Analysis |
|---|---|---|---|---|---|---|---|---|---|
| AfontovaGora3_d | Russia_LUP | Russia | Palaeolithic | 17930 | F | 56.02 | 92.87 | Fu et al. (2016) doi: 10.1038/nature17993 | PCA, Admixture |
| AH1.SG | Zagros_N | Iran | Neolithic | 9900 | F | 34.19 | 48.37 | Broushaki et al. (2016) doi: 10.1126/science.aaf7943 | PCA, Admixture |
| AH2.SG | Zagros_N | Iran | Neolithic | 9931 | M | 34.19 | 48.37 | Broushaki et al. (2016) doi: 10.1126/science.aaf7943 | PCA, Admixture |
| AH4.SG | Zagros_N | Iran | Neolithic | 9930 | F | 34.19 | 48.37 | Broushaki et al. (2016) doi: 10.1126/science.aaf7943 | PCA, Admixture, |
| ANI159-ANI181 | Balkans_N | Bulgaria | Neolithic | 6571 | M | 43.21 | 27.86 | Mathieson et al. (2018) doi: 10.1038/nature25778 | PCA, Admixture, |
| ANI160 | Balkans_N | Bulgaria | Neolithic | 6550 | M | 43.21 | 27.86 | Mathieson et al. (2018) doi: 10.1038/nature25778 | PCA, Admixture, |
| ANI163 | Balkans_N | Bulgaria | Neolithic | 6577 | F | 43.21 | 27.86 | Mathieson et al. (2018) doi: 10.1038/nature25778 | PCA, Admixture, |
| atp002.SG | SWEurope_C | Spain | Copper Age | 4740 | M | 42.35 | -3.52 | Valdiosera et al. (2018) doi: 10.1073/pnas.1717762115 | PCA, Admixture, |
| atp005.SG | SWEurope_N | Spain | Neolithic | 7074 | M | 42.35 | -3.52 | Valdiosera et al. (2018) doi: 10.1073/pnas.1717762115 | PCA, Admixture, |
| atp12-1420.SG | SWEurope_C | Spain | Copper Age | 4896 | M | 42.35 | -3.52 | Valdiosera et al. (2018) doi: 10.1073/pnas.1717762115 | PCA, Admixture, |
| baa001.SG | SAfrica_LSA | South Africa | Palaeolithic | 1909 | M | 29.32 | 31.13 | Schlebusch et al. (2017) doi: 10.1126/science.aao6266 | PCA, Admixture, |
| bab001.SG | SAfrica_LSA | South Africa | Palaeolithic | 2041 | M | 29.32 | 31.13 | Schlebusch et al. (2017) doi: 10.1126/science.aao6266 | PCA, Admixture, |
| Bar31.SG | Anatolia_N | Turkey | Neolithic | 8279 | M | 40.30 | 29.57 | Hofmanova et al. (2016) doi: 10.1073/pnas.1523951113 | PCA, Admixture, |
| Bar8.SG | Anatolia_N | Turkey | Neolithic | 8071 | F | 40.30 | 29.57 | Hofmanova et al. (2016) doi: 10.1073/pnas.1523951113 | PCA, Admixture, NNLS, |
| Bichon.SG | CEurope_HG | Switzerland | Palaeolithic | 13665 | M | 47.10 | 6.87 | Jones et al. (2015) doi: 10.1038/ncomms9912 | PCA, Admixture, NNLS, Right-pop in qpWave and qpAdm (Outgroup) |
| Bon001.SG | Anatolia_N | Turkey | Neolithic | 10032 | M | 37.75 | 32.86 | Kilinc et al. (2016) doi: 10.1016/j.cub.2016.07.057 | PCA, Admixture, |
| Bon002.SG | Anatolia_N | Turkey | Neolithic | 10078 | F | 37.75 | 32.86 | Kilinc et al. (2016) doi: 10.1016/j.cub.2016.07.057 | PCA, Admixture, |
| Bon004.SG | Anatolia_N | Turkey | Neolithic | 10076 | M | 37.75 | 32.86 | Kilinc et al. (2016) doi: 10.1016/j.cub.2016.07.057 | PCA, Admixture, |
| BOT14.SG | CAsia_Botai_EN | Kazakhstan | Neolithic | 5264 | M | 53.31 | 67.65 | Damgaard et al. (2018) doi: 10.1126/science.aar7711 | PCA, Admixture, |
| BOT15.SG | CAsia_Botai_EN | Kazakhstan | Neolithic | 5135 | M | 53.31 | 67.65 | Damgaard et al. (2018) doi: 10.1126/science.aar7711 | PCA, Admixture, |
| BOT2016.SG | CAsia_Botai_EN | Kazakhstan | Neolithic | 5450 | F | 53.31 | 67.65 | Damgaard et al. (2018) doi: 10.1126/science.aar7711 | PCA, Admixture, |
| c40331.SG | SWEurope_N | Spain | Neolithic | 5650 | M | 37.36 | -4.25 | Valdiosera et al. (2018) doi: 10.1073/pnas.1717762115 | PCA, Admixture, |
| CabecoArruda117B.SG | SWEurope_N | Portugal | Neolithic | 5050 | M | 39.11 | -8.66 | Martiniano et al. (2017) doi: 10.1371/journal.pgen.1006852 | PCA, Admixture, |

| | | | | | | | | | |
|---|---|---|---|---|---|---|---|---|---|
| CabecoArruda122A.SG | SWEurope_N | Portugal | Neolithic | 5050 | M | 39.11 | -8.66 | Martiniano et al. (2017) doi: 10.1371/journal.pgen.1006852 | PCA, Admixture, |
| CB13.SG | SWEurope_N | Spain | Neolithic | 7348 | F | 41.37 | 1.89 | Olalde et al. (2015) doi: 10.1093/molbev/msv181 | PCA, Admixture, |
| cha001.SG | SAfrica_LSA_IA | South Africa | Iron Age | 365 | F | 29.09 | 29.33 | Schlebusch et al. (2017) doi: 10.1126/science.aao6266 | PCA, Admixture, |
| CovaMoura364.SG | SWEurope_N | Portugal | Neolithic | 4900 | M | 38.75 | -9.22 | Martiniano et al. (2017) doi: 10.1371/journal.pgen.1006852 | PCA, Admixture, |
| CovaMoura9B.SG | SWEurope_N | Portugal | Neolithic | 4900 | F | 38.75 | -9.22 | Martiniano et al. (2017) doi: 10.1371/journal.pgen.1006852 | PCA, Admixture, |
| DA38.SG | CAsia_XiongNu_WE | Mongolia | Historical or Others | 2152 | F | 49.27 | 101.72 | Damgaard et al. (2018) doi:10.1038/s41586-018-0094-2 | Right-pop in qpWave and qpAdm (Outgroup), |
| DA39.SG | CAsia_XiongNu | Mongolia | Historical or Others | 1990 | M | 48.02 | 101.35 | Damgaard et al. (2018) doi:10.1038/s41586-018-0094-2 | Right-pop in qpWave and qpAdm (Outgroup), |
| DA43.SG | CAsia_XiongNu | Mongolia | Historical or Others | 2100 | M | 42.53 | 105.18 | Damgaard et al. (2018) doi:10.1038/s41586-018-0094-2 | Right-pop in qpWave and qpAdm (Outgroup), |
| DA45.SG | CAsia_XiongNu | Mongolia | Historical or Others | 2062 | M | 42.53 | 105.18 | Damgaard et al. (2018) doi:10.1038/s41586-018-0094-2 | Right-pop in qpWave and qpAdm (Outgroup), |
| DolmenAnsiao96B.SG | SWEurope_N | Portugal | Neolithic | 5450 | M | 39.91 | -8.44 | Martiniano et al. (2017) doi: 10.1371/journal.pgen.1006852 | PCA, Admixture, |
| ela001.SG | SAfrica_LSA_IA | South Africa | Iron Age | 493 | F | 32.32 | 18.32 | Schlebusch et al. (2017) doi: 10.1126/science.aao6266 | PCA, Admixture, |
| ElMiron_d | SWEurope_LUP | Spain | Palaeolithic | 18720 | F | 43.26 | -3.45 | Fu et al. (2016) doi: 10.1038/nature17993 | PCA, Admixture, NNLS, |
| GoyetQ116-1_published | WEurope_EUP | Belgium | Palaeolithic | 34795 | M | 50.26 | 4.28 | Fu et al. (2016) doi: 10.1038/nature17993 | PCA, Admixture, |
| HohleFels49_published_d | CEurope_LUP | Germany | Palaeolithic | 15130 | M | 48.22 | 9.45 | Fu et al. (2016) doi: 10.1038/nature17993 | PCA, Admixture, |
| I0017.SG | NEurope_HG | Sweden | Palaeolithic | 7626 | M | 58.54 | 15.05 | Lazaridis et al. (2014) doi: 10.1038/nature13673 | PCA, Admixture, |
| I0061 | NWRussia_HG | Russia | Palaeolithic | 8280 | M | 61.65 | 35.65 | Mathieson et al. (2015) doi: 10.1038/nature16152 (1240k of same sample with 390k in Haak, Lazaridis, et al. (2015) doi: 10.1038/nature14317) | PCA, Admixture, Left-pop in qpWave and qpAdm for Basal Eurasia, |
| I0070 | SEEurope_Minoan_BA | Greece | Bronze Age | 4000 | M | 35.08 | 25.83 | Lazaridis et al. (2017) doi: 10.1038/nature23310 | PCA, Admixture, |
| I0071 | SEEurope_Minoan_BA | Greece | Bronze Age | 4000 | F | 35.08 | 25.83 | Lazaridis et al. (2017) doi: 10.1038/nature23310 | PCA, Admixture, |
| I0073 | SEEurope_Minoan_BA | Greece | Bronze Age | 4000 | M | 35.08 | 25.83 | Lazaridis et al. (2017) doi: 10.1038/nature23310 | PCA, Admixture, |
| I0074 | SEEurope_Minoan_BA | Greece | Bronze Age | 4000 | F | 35.08 | 25.83 | Lazaridis et al. (2017) doi: 10.1038/nature23310 | PCA, Admixture, |
| I0100 | CEurope_LBK_EN | Germany | Neolithic | 6977 | F | 51.90 | 11.05 | Lipson et al. (2017) doi: 10.1038/nature24476 (supplement of Mathieson et al. (2015) doi: 10.1038/nature16152, which itself was supplement of Haak, Lazaridis, et al. (2015) doi: 10.1038/nature14317) | PCA, Admixture, Right-pop in qpWave and qpAdm (Outgroup), |
| I0103 | CEurope_LN | Germany | Neolithic | 4473 | F | 51.42 | 11.68 | Mathieson et al. (2015) doi: 10.1038/nature16152 (1240k of same sample with 390k in Haak, Lazaridis, et al. (2015) doi: 10.1038/nature14317) | PCA, Admixture, Right-pop in qpWave and qpAdm (Outgroup), |
| I0112 | CEurope_BellBeaker_BA | Germany | Bronze Age | 4250 | F | 51.79 | 11.14 | Mathieson et al. (2015) doi: 10.1038/nature16152 (1240k of same sample with 390k in Haak, Lazaridis, et al. (2015) doi: 10.1038/nature14317) | PCA, Admixture, NNLS, Right-pop in qpWave and qpAdm (Outgroup), |
| I0231_published | RussiaSteppe_EMBA | Russia | Bronze Age | 4800 | M | 52.71 | 49.47 | Mathieson et al. (2015) doi: 10.1038/nature16152 (1240k of same with 390k in Haak, Lazaridis, et al. (2015) doi: 10.1038/nature14317) | PCA, Admixture, NNLS, |
| I0247 | RussiaSteppe_IA | Russia | Iron Age | 2239 | M | 52.43 | 51.16 | Mathieson et al. (2015) doi: 10.1038/nature16152 | PCA, Admixture, |
| I0405 | SWEurope_N | Spain | Neolithic | 5700 | M | 41.25 | -2.33 | Mathieson et al. (2015) doi: 10.1038/nature16152 (1240k of same sample with 390k in Haak, Lazaridis, et al. (2015) doi: | PCA, Admixture, |

| | | | | | | | | | |
|---|---|---|---|---|---|---|---|---|---|
| | | | | | | | | 10.1038/nature14317) | |
| I0406 | SWEurope_N | Spain | Neolithic | 5700 | M | 41.25 | -2.33 | Mathieson et al. (2015) doi: 10.1038/nature16152 (1240k of same sample with 390k in Haak, Lazaridis, et al. (2015) doi: 10.1038/nature14317) | PCA, Admixture, |
| I0407 | SWEurope_N | Spain | Neolithic | 5700 | F | 41.25 | -2.33 | Mathieson et al. (2015) doi: 10.1038/nature16152 (1240k of samesame sample with 390k in Haak, Lazaridis, et al. (2015) doi: 10.1038/nature14317) | PCA, Admixture, |
| I0408 | SWEurope_N | Spain | Neolithic | 5727 | F | 41.25 | -2.33 | Mathieson et al. (2015) doi: 10.1038/nature16152 (1240k of same sample with 390k in Haak, Lazaridis, et al. (2015) doi: 10.1038/nature14317) | PCA, Admixture, |
| I0409 | SWEurope_N | Spain | Neolithic | 7215 | F | 42.50 | 0.50 | Mathieson et al. (2015) doi: 10.1038/nature16152 (1240k of same sample with 390k in Haak, Lazaridis, et al. (2015) doi: 10.1038/nature14317) | PCA, Admixture, |
| I0410 | SWEurope_N | Spain | Neolithic | 7130 | M | 42.50 | 0.50 | Mathieson et al. (2015) doi: 10.1038/nature16152 (1240k of same sample with 390k in Haak, Lazaridis, et al. (2015) doi: 10.1038/nature14317) | PCA, Admixture, |
| I0411 | SWEurope_N | Spain | Neolithic | 7131 | M | 42.50 | 0.50 | Mathieson et al. (2015) doi: 10.1038/nature16152 (1240k of same sample with 390k in Haak, Lazaridis, et al. (2015) doi: 10.1038/nature14317) | PCA, Admixture, |
| I0412 | SWEurope_N | Spain | Neolithic | 7144 | M | 42.50 | 0.50 | Mathieson et al. (2015) doi: 10.1038/nature16152 (1240k of same with 390k in Haak, Lazaridis, et al. (2015) doi: 10.1038/nature14317) | PCA, Admixture, |
| I0413 | SWEurope_N | Spain | Neolithic | 7139 | F | 42.50 | 0.50 | Mathieson et al. (2015) doi: 10.1038/nature16152 (1240k of same sample with 390k in Haak, Lazaridis, et al. (2015) doi: 10.1038/nature14317) | PCA, Admixture, |
| I0443 | RussiaSteppe_EMBA | Russia | Bronze Age | 4950 | M | 53.38 | 50.39 | Mathieson et al. (2015) doi: 10.1038/nature16152 (1240k of same sample with 390k in Haak, Lazaridis, et al. (2015) doi: 10.1038/nature14317) | PCA, Admixture, |
| I0585 | SWEurope_HG | Spain | Palaeolithic | 7815 | M | 42.91 | -5.38 | Mathieson et al. (2015) doi: 10.1038/nature16152 (1240k of same sample with shotgun in Olalde et al. (2014) doi: 10.1038/nature12960) | PCA, Admixture, NNLS, |
| I0589_all | SEAfrica_MIA | Tanzania | Iron Age | 1335 | F | -6.40 | 39.50 | Skoglund et al. (2017) doi: 10.1016/j.cell.2017.08.049 | PCA, Admixture, |
| I0595 | EAfrica_MIA_LIA | Kenya | Iron Age | 410 | M | -3.70 | 39.70 | Skoglund et al. (2017) doi: 10.1016/j.cell.2017.08.049 | PCA, Admixture, |
| I0626_all | SEAsia_N | Vietnam | Neolithic | 3750 | M | 20.13 | 105.98 | Lipson et al. (2018) doi: 10.1126/science.aat3188 | PCA, Admixture, |
| I0627_all | SEAsia_N | Vietnam | Neolithic | 3964 | F | 20.13 | 105.98 | Lipson et al. (2018) doi: 10.1126/science.aat3188 | PCA, Admixture, |
| I0676 | Balkans_N | Macedonia | Neolithic | 7807 | M | 41.90 | 21.35 | Mathieson et al. (2018) doi: 10.1038/nature25778 | PCA, Admixture, |
| I0679_d | Balkans_N | Bulgaria | Neolithic | 7622 | F | 42.02 | 25.60 | Mathieson et al. (2018) doi: 10.1038/nature25778 | PCA, Admixture, |
| I0698_published | Balkans_N | Bulgaria | Neolithic | 7900 | M | 42.10 | 25.75 | Mathieson et al. (2018) doi: 10.1038/nature25778 | PCA, Admixture, |
| I0700 | Balkans_N | Bulgaria | Neolithic | 7934 | M | 43.98 | 26.40 | Mathieson et al. (2018) doi: 10.1038/nature25778 | PCA, Admixture, |
| I0704_published | Balkans_N | Bulgaria | Neolithic | 7889 | F | 43.16 | 25.88 | Mathieson et al. (2018) doi: 10.1038/nature25778 | PCA, Admixture, |
| I0706 | Balkans_N | Bulgaria | Neolithic | 7905 | M | 43.16 | 25.88 | Mathieson et al. (2018) doi: 10.1038/nature25778 | PCA, Admixture, |
| I0707 | Anatolia_N | Turkey | Neolithic | 8092 | F | 40.30 | 29.57 | Mathieson et al. (2015) doi: 10.1038/nature16152 | PCA, Admixture, |
| I0708 | Anatolia_N | Turkey | Neolithic | 8097 | M | 40.30 | 29.57 | Mathieson et al. (2015) doi: 10.1038/nature16152 | PCA, Admixture, |
| I0709 | Anatolia_N | Turkey | Neolithic | 8086 | M | 40.30 | 29.57 | Mathieson et al. (2015) doi: 10.1038/nature16152 | PCA, Admixture, |

| | | | | | | | | | |
|---|---|---|---|---|---|---|---|---|---|
| I0723 | Anatolia_N | Turkey | Neolithic | 7870 | M | 40.26 | 29.65 | Mathieson et al. (2015) doi: 10.1038/nature16152 | PCA, Admixture, |
| I0724 | Anatolia_N | Turkey | Neolithic | 7950 | M | 40.26 | 29.65 | Mathieson et al. (2015) doi: 10.1038/nature16152 | PCA, Admixture, |
| I0725 | Anatolia_N | Turkey | Neolithic | 7950 | F | 40.26 | 29.65 | Mathieson et al. (2015) doi: 10.1038/nature16152 | PCA, Admixture, |
| I0726 | Anatolia_N | Turkey | Neolithic | 7950 | F | 40.26 | 29.65 | Mathieson et al. (2015) doi: 10.1038/nature16152 | PCA, Admixture, |
| I0727 | Anatolia_N | Turkey | Neolithic | 7950 | M | 40.26 | 29.65 | Mathieson et al. (2015) doi: 10.1038/nature16152 | PCA, Admixture, |
| I0736 | Anatolia_N | Turkey | Neolithic | 8300 | F | 40.30 | 29.57 | Mathieson et al. (2015) doi: 10.1038/nature16152 | PCA, Admixture, |
| I0744 | Anatolia_N | Turkey | Neolithic | 8273 | M | 40.30 | 29.57 | Mathieson et al. (2015) doi: 10.1038/nature16152 | PCA, Admixture, |
| I0745 | Anatolia_N | Turkey | Neolithic | 8251 | M | 40.30 | 29.57 | Mathieson et al. (2015) doi: 10.1038/nature16152 | PCA, Admixture, |
| I0746 | Anatolia_N | Turkey | Neolithic | 7930 | M | 40.30 | 29.57 | Mathieson et al. (2015) doi: 10.1038/nature16152 | PCA, Admixture, |
| I0854 | Anatolia_N | Turkey | Neolithic | 8106 | F | 40.30 | 29.57 | Mathieson et al. (2015) doi: 10.1038/nature16152 | PCA, Admixture, |
| I0861 | Natufian | Israel | Palaeolithic | 12750 | M | 32.65 | 35.07 | Lazaridis et al. (2016) doi: 10.1038/nature19310 | PCA, Admixture, |
| I0867 | Levant_PPNB | Israel | Neolithic | 8700 | M | 31.79 | 35.17 | Lazaridis et al. (2016) doi: 10.1038/nature19310 | PCA, Admixture, NNLS, Left-pop in qpWave and qpAdm, |
| I1048 | SEAfrica_LSA | Tanzania | Palaeolithic | 1358 | F | -4.90 | 39.60 | Skoglund et al. (2017) doi: 10.1016/j.cell.2017.08.049 | PCA, Admixture, |
| I1072 | Natufian | Israel | Palaeolithic | 12750 | M | 32.65 | 35.07 | Lazaridis et al. (2016) doi: 10.1038/nature19310 | PCA, Admixture, NNLS, Left-pop in qpWave and qpAdm, Left-pop in qpWave and qpAdm for Basal Eurasia, *f4*-statistics, *f4*-statistics for Neanderthal |
| I1096 | Anatolia_N | Turkey | Neolithic | 8300 | M | 40.30 | 29.57 | Mathieson et al. (2015) doi: 10.1038/nature16152 | PCA, Admixture, |
| I1097 | Anatolia_N | Turkey | Neolithic | 8288 | M | 40.30 | 29.57 | Mathieson et al. (2015) doi: 10.1038/nature16152 | PCA, Admixture, |
| I1098 | Anatolia_N | Turkey | Neolithic | 8288 | F | 40.30 | 29.57 | Mathieson et al. (2015) doi: 10.1038/nature16152 | PCA, Admixture, |
| I1099 | Anatolia_N | Turkey | Neolithic | 8300 | M | 40.30 | 29.57 | Mathieson et al. (2015) doi: 10.1038/nature16152 | PCA, Admixture, |
| I1100 | Anatolia_N | Turkey | Neolithic | 8300 | F | 40.30 | 29.57 | Mathieson et al. (2015) doi: 10.1038/nature16152 | PCA, Admixture, |
| I1101 | Anatolia_N | Turkey | Neolithic | 8300 | M | 40.30 | 29.57 | Mathieson et al. (2015) doi: 10.1038/nature16152 | PCA, Admixture, |
| I1102 | Anatolia_N | Turkey | Neolithic | 8300 | M | 40.30 | 29.57 | Mathieson et al. (2015) doi: 10.1038/nature16152 | PCA, Admixture, |
| I1103 | Anatolia_N | Turkey | Neolithic | 8300 | M | 40.30 | 29.57 | Mathieson et al. (2015) doi: 10.1038/nature16152 | PCA, Admixture, |
| I1108 | Balkans_N | Bulgaria | Neolithic | 7875 | M | 43.98 | 26.40 | Mathieson et al. (2018) doi: 10.1038/nature25778 | PCA, Admixture, |
| I1109 | Balkans_N | Bulgaria | Neolithic | 7635 | F | 43.98 | 26.40 | Mathieson et al. (2018) doi: 10.1038/nature25778 | PCA, Admixture, |
| I1113 | Balkans_N | Bulgaria | Neolithic | 7983 | F | 43.98 | 26.40 | Mathieson et al. (2018) doi: 10.1038/nature25778 | PCA, Admixture, |
| I1137_all | SEAsia_N | Vietnam | Neolithic | 3764 | M | 20.13 | 105.98 | Lipson et al. (2018) doi: 10.1126/science.aat3188 | PCA, Admixture, |
| I1290 | Zagros_N | Iran | Neolithic | 9846 | F | 34.45 | 48.12 | Lazaridis et al. (2016) doi: 10.1038/nature19310 | PCA, Admixture, |
| I1293_all | Iran_EpiPalaeolithic | Iran | Palaeolithic | 10800 | M | 35.59 | 53.50 | Lazaridis et al. (2016) doi: 10.1038/nature19310 | PCA, Admixture, |

| | | | | | | | | | |
|---|---|---|---|---|---|---|---|---|---|
| I1295 | Balkans_N | Bulgaria | Neolithic | 7752 | M | 43.98 | 26.40 | Mathieson et al. (2018) doi: 10.1038/nature25778 | PCA, Admixture, |
| I1296_published | Balkans_N | Bulgaria | Neolithic | 7675 | M | 43.98 | 26.40 | Mathieson et al. (2018) doi: 10.1038/nature25778 | PCA, Admixture, |
| I1297 | Balkans_N | Bulgaria | Neolithic | 7817 | F | 43.98 | 26.40 | Mathieson et al. (2018) doi: 10.1038/nature25778 | PCA, Admixture, |
| I1407 | Caucasus_C | Armenia | Copper Age | 5875 | M | 39.73 | 45.20 | Lazaridis et al. (2016) doi: 10.1038/nature19310 | PCA, Admixture, |
| I1414_published | Levant_PPNB | Jordan | Neolithic | 10050 | M | 31.99 | 35.98 | Lazaridis et al. (2016) doi: 10.1038/nature19310 | PCA, Admixture, |
| I1415_published | Levant_PPNB | Jordan | Neolithic | 9875 | M | 31.99 | 35.98 | Lazaridis et al. (2016) doi: 10.1038/nature19310 | PCA, Admixture, |
| I1499 | CEurope_MN | Hungary | Neolithic | 7060 | F | 48.52 | 21.17 | Mathieson et al. (2015) doi: 10.1038/nature16152 (capture of same sample shotgunned in Gamba et al. (2014) doi: 10.1038/ncomms6257) | PCA, Admixture, Right-pop in qpWave and qpAdm (Outgroup), |
| I1580 | Anatolia_N | Turkey | Neolithic | 8195 | F | 40.30 | 29.57 | Mathieson et al. (2015) doi: 10.1038/nature16152 | PCA, Admixture, |
| I1581 | Anatolia_N | Turkey | Neolithic | 8254 | F | 40.30 | 29.57 | Mathieson et al. (2015) doi: 10.1038/nature16152 | PCA, Admixture, |
| I1583 | Anatolia_N | Turkey | Neolithic | 8281 | M | 40.30 | 29.57 | Mathieson et al. (2015) doi: 10.1038/nature16152 | PCA, Admixture, |
| I1584 | Anatolia_C | Turkey | Copper Age | 5776 | F | 40.30 | 29.57 | Lazaridis et al. (2016) doi: 10.1038/nature19310 | PCA, Admixture, |
| I1633 | Caucasus_EBA | Armenia | Bronze Age | 4465 | F | 40.65 | 45.12 | Lazaridis et al. (2016) doi: 10.1038/nature19310 | PCA, Admixture, NNLS, Left-pop in qpWave and qpAdm, |
| I1656 | Caucasus_MBA | Armenia | Bronze Age | 3402 | F | 40.38 | 43.94 | Lazaridis et al. (2016) doi: 10.1038/nature19310 | PCA, Admixture, NNLS, |
| I1658 | Caucasus_EBA | Armenia | Bronze Age | 5170 | F | 40.39 | 43.89 | Lazaridis et al. (2016) doi: 10.1038/nature19310 | PCA, Admixture, |
| I1671 | Zagros_N | Iran | Neolithic | 7698 | M | 34.50 | 47.96 | Lazaridis et al. (2016) doi: 10.1038/nature19310 | PCA, Admixture, |
| I1679 | Levant_PPNC | Jordan | Neolithic | 8800 | F | 31.99 | 35.98 | Lazaridis et al. (2016) doi: 10.1038/nature19310 | PCA, Admixture, |
| I1680 | SEAsia_IA | Cambodia | Iron Age | 1810 | M | 10.99 | 104.98 | Lipson et al. (2018) doi: 10.1126/science.aat3188 | PCA, Admixture, |
| I1685_published | Natufian | Israel | Palaeolithic | 12750 | M | 32.65 | 35.07 | Lazaridis et al. (2016) doi: 10.1038/nature19310 | PCA, Admixture, |
| I1699 | Levant_PPNC | Jordan | Neolithic | 8700 | F | 31.99 | 35.98 | Lazaridis et al. (2016) doi: 10.1038/nature19310 | PCA, Admixture, |
| I1700_published | Levant_PPNB | Jordan | Neolithic | 10050 | M | 31.99 | 35.98 | Lazaridis et al. (2016) doi: 10.1038/nature19310 | PCA, Admixture, |
| I1704 | Levant_PPNB | Jordan | Neolithic | 9202 | F | 31.99 | 35.98 | Lazaridis et al. (2016) doi: 10.1038/nature19310 | PCA, Admixture, |
| I1705 | Levant_EBA | Jordan | Bronze Age | 4032 | M | 31.99 | 35.98 | Lazaridis et al. (2016) doi: 10.1038/nature19310 | PCA, Admixture, NNLS, Left-pop in qpWave and qpAdm, |
| I1706 | Levant_EBA | Jordan | Bronze Age | 4345 | F | 31.99 | 35.98 | Lazaridis et al. (2016) doi: 10.1038/nature19310 | PCA, Admixture, |
| I1707 | Levant_PPNB | Jordan | Neolithic | 9582 | M | 31.99 | 35.98 | Lazaridis et al. (2016) doi: 10.1038/nature19310 | PCA, Admixture, |
| I1710 | Levant_PPNB | Jordan | Neolithic | 9580 | M | 31.99 | 35.98 | Lazaridis et al. (2016) doi: 10.1038/nature19310 | PCA, Admixture, |
| I1727 | Levant_PPNB | Jordan | Neolithic | 10050 | M | 31.99 | 35.98 | Lazaridis et al. (2016) doi: 10.1038/nature19310 | PCA, Admixture, |
| I1730 | Levant_EBA | Jordan | Bronze Age | 4344 | M | 31.99 | 35.98 | Lazaridis et al. (2016) doi: 10.1038/nature19310 | PCA, Admixture, |
| I1859_all | SEAsia_N | Vietnam | Neolithic | 3766 | F | 20.13 | 105.98 | Lipson et al. (2018) doi: 10.1126/science.aat3188 | PCA, Admixture, |

| | | | | | | | | | |
|---|---|---|---|---|---|---|---|---|---|
| I1945_published | Zagros_N | Iran | Neolithic | 9800 | M | 34.45 | 48.12 | Lazaridis et al. (2016) doi: 10.1038/nature19310 | PCA, Admixture, |
| I1949_published | Zagros_N | Iran | Neolithic | 10052 | M | 34.45 | 48.12 | Lazaridis et al. (2016) doi: 10.1038/nature19310 | PCA, Admixture, |
| I1972 | SWEurope_N | Spain | Neolithic | 6710 | F | 42.63 | -3.12 | Lipson et al. (2017) doi: 10.1038/nature24476 | PCA, Admixture, |
| I1979 | SEurope_BellBeaker_BA | Italy | Bronze Age | 4015 | F | 44.78 | 10.29 | Olalde et al. (2018) doi: 10.1038/nature25738 | PCA, Admixture, |
| I2158_published | SEurope_HG | Italy | Palaeolithic | 14275 | F | 37.93 | 12.33 | Mathieson et al. (2018) doi: 10.1038/nature25778 | PCA, Admixture, |
| I2199 | SWEurope_N | Spain | Neolithic | 7074 | F | 42.63 | -3.12 | Lipson et al. (2017) doi: 10.1038/nature24476 | PCA, Admixture, |
| I2215_published | Balkans_N | Bulgaria | Neolithic | 7980 | M | 43.98 | 26.40 | Mathieson et al. (2018) doi: 10.1038/nature25778 | PCA, Admixture, |
| I2216_published | Balkans_N | Bulgaria | Neolithic | 7845 | F | 43.98 | 26.40 | Mathieson et al. (2018) doi: 10.1038/nature25778 | PCA, Admixture, |
| I2318 | SEEurope_Peloponnese_N | Greece | Neolithic | 5945 | F | 37.42 | 23.13 | Mathieson et al. (2018) doi: 10.1038/nature25778 | PCA, Admixture, |
| I2477 | SEurope_BellBeaker_BA | Italy | Bronze Age | 4015 | F | 44.78 | 10.29 | Olalde et al. (2018) doi: 10.1038/nature25738 | PCA, Admixture, NNLS, |
| I2478 | SEurope_BellBeaker_BA | Italy | Bronze Age | 4018 | M | 44.78 | 10.29 | Olalde et al. (2018) doi: 10.1038/nature25738 | PCA, Admixture, |
| I2495 | Anatolia_EBA | Turkey | Bronze Age | 4377 | M | 37.92 | 30.71 | Lazaridis et al. (2017) doi: 10.1038/nature23310 | PCA, Admixture, NNLS, |
| I2497_all | SEAsia_BA | Vietnam | Bronze Age | 2000 | F | 19.80 | 105.80 | Lipson et al. (2018) doi: 10.1126/science.aat3188 | PCA, Admixture |
| I2499 | Anatolia_EBA | Turkey | Bronze Age | 4604 | F | 37.92 | 30.71 | Lazaridis et al. (2017) doi: 10.1038/nature23310 | PCA, Admixture |
| I2521 | Balkans_N | Bulgaria | Neolithic | 7505 | M | 43.16 | 25.88 | Mathieson et al. (2018) doi: 10.1038/nature25778 | PCA, Admixture |
| I2526 | Balkans_N | Bulgaria | Neolithic | 7336 | F | 43.14 | 25.61 | Mathieson et al. (2018) doi: 10.1038/nature25778 | PCA, Admixture |
| I2529_published | Balkans_N | Bulgaria | Neolithic | 7601 | M | 42.10 | 25.75 | Mathieson et al. (2018) doi: 10.1038/nature25778 | PCA, Admixture |
| I2683 | Anatolia_EBA | Turkey | Bronze Age | 4494 | F | 37.92 | 30.71 | Lazaridis et al. (2017) doi: 10.1038/nature23310 | PCA, Admixture, |
| I2731_all | SEAsia_N | Vietnam | Neolithic | 3717 | M | 20.13 | 105.98 | Lipson et al. (2018) doi: 10.1126/science.aat3188 | PCA, Admixture, |
| I2947 | SEAsia_N | Vietnam | Neolithic | 3750 | F | 20.13 | 105.98 | Lipson et al. (2018) doi: 10.1126/science.aat3188 | PCA, Admixture, |
| I2948_all | SEAsia_BA | Vietnam | Bronze Age | 1948 | M | 19.80 | 105.80 | Lipson et al. (2018) doi: 10.1126/science.aat3188 | PCA, Admixture, |
| I2966_all | SEAfrica_HG | Malawi | Palaeolithic | 7950 | M | -11.66 | 33.64 | Skoglund et al. (2017) doi: 10.1016/j.cell.2017.08.049 | PCA, Admixture, |
| I2967_all | SEAfrica_HG | Malawi | Palaeolithic | 8065 | F | -11.66 | 33.64 | Skoglund et al. (2017) doi: 10.1016/j.cell.2017.08.049 | PCA, Admixture, |
| I3433 | Balkans_N | Croatia | Neolithic | 7814 | F | 43.59 | 16.65 | Mathieson et al. (2018) doi: 10.1038/nature25778 | PCA, Admixture, |
| I3498 | Balkans_N | Croatia | Neolithic | 7698 | M | 45.34 | 18.70 | Mathieson et al. (2018) doi: 10.1038/nature25778 | PCA, Admixture, |
| I3499 | Balkans_N | Croatia | Neolithic | 4725 | M | 45.34 | 18.70 | Mathieson et al. (2018) doi: 10.1038/nature25778 | PCA, Admixture, |
| I3708 | SEEurope_Peloponnese_N | Greece | Neolithic | 6550 | F | 36.64 | 22.38 | Mathieson et al. (2018) doi: 10.1038/nature25778 | PCA, Admixture, |
| I3709 | SEEurope_Peloponnese_N | Greece | Neolithic | 5847 | F | 36.64 | 22.38 | Mathieson et al. (2018) doi: 10.1038/nature25778 | PCA, Admixture, |
| I3726_all | SEafrica_N | Tanzania | Neolithic | 3079 | F | -4.26 | 35.32 | Skoglund et al. (2017) doi: 10.1016/j.cell.2017.08.049 | PCA, Admixture, |

| | | | | | | | | | |
|---|---|---|---|---|---|---|---|---|---|
| I3879 | Balkans_N | Bulgaria | Neolithic | 7828 | M | 43.98 | 26.40 | Mathieson et al. (2018) doi: 10.1038/nature25778 | PCA, Admixture, |
| I3920 | SEEurope_Peloponnese_N | Greece | Neolithic | 5770 | F | 36.64 | 22.38 | Mathieson et al. (2018) doi: 10.1038/nature25778 | PCA, Admixture, |
| I3947 | Balkans_N | Croatia | Neolithic | 7836 | M | 43.59 | 16.65 | Mathieson et al. (2018) doi: 10.1038/nature25778 | PCA, Admixture, |
| I3948 | Balkans_N | Croatia | Neolithic | 7860 | M | 43.59 | 16.65 | Mathieson et al. (2018) doi: 10.1038/nature25778 | PCA, Admixture, |
| I4167 | Balkans_N | Croatia | Neolithic | 6624 | M | 45.34 | 18.70 | Mathieson et al. (2018) doi: 10.1038/nature25778 | PCA, Admixture, |
| I4168 | Balkans_N | Croatia | Neolithic | 6600 | F | 45.34 | 18.70 | Mathieson et al. (2018) doi: 10.1038/nature25778 | PCA, Admixture, |
| I4421_all_published | SEAfrica_LSA | Malawi | Palaeolithic | 5200 | F | -14.38 | 33.82 | Skoglund et al. (2017) doi: 10.1016/j.cell.2017.08.049 | PCA, Admixture, |
| I4426_all_published | SEAfrica_LSA | Malawi | Palaeolithic | 2517 | F | -10.79 | 33.77 | Skoglund et al. (2017) doi: 10.1016/j.cell.2017.08.049 | PCA, Admixture, |
| I4427_all_published | SEAfrica_LSA | Malawi | Palaeolithic | 6061 | M | -10.79 | 33.77 | Skoglund et al. (2017) doi: 10.1016/j.cell.2017.08.049 | PCA, Admixture, |
| I4468_all_published | SEAfrica_LSA | Malawi | Palaeolithic | 6087 | M | -10.79 | 33.77 | Skoglund et al. (2017) doi: 10.1016/j.cell.2017.08.049 | PCA, Admixture, |
| I4930 | SEurope_C | Italy | Copper Age | 4721 | F | 37.73 | 12.96 | Olalde et al. (2018) doi: 10.1038/nature25738 | PCA, Admixture, |
| I5071 | Balkans_N | Croatia | Neolithic | 7571 | F | 45.37 | 13.56 | Mathieson et al. (2018) doi: 10.1038/nature25778 | PCA, Admixture, |
| I5072 | Balkans_N | Croatia | Neolithic | 7551 | M | 45.37 | 13.56 | Mathieson et al. (2018) doi: 10.1038/nature25778 | PCA, Admixture, |
| I5077 | Balkans_N | Croatia | Neolithic | 7026 | M | 45.55 | 18.75 | Mathieson et al. (2018) doi: 10.1038/nature25778 | PCA, Admixture, |
| I5078 | Balkans_N | Croatia | Neolithic | 6569 | M | 45.55 | 18.75 | Mathieson et al. (2018) doi: 10.1038/nature25778 | PCA, Admixture, NNLS, |
| I5079 | Balkans_N | Croatia | Neolithic | 5485 | F | 45.49 | 17.64 | Mathieson et al. (2018) doi: 10.1038/nature25778 | PCA, Admixture, |
| I5235 | Balkans_HG | Serbia | Palaeolithic | 10835 | M | 44.60 | 22.01 | Mathieson et al. (2018) doi: 10.1038/nature25778 | PCA, Admixture, |
| I5236 | Balkans_HG | Serbia | Palaeolithic | 10008 | M | 44.60 | 22.01 | Mathieson et al. (2018) doi: 10.1038/nature25778 | PCA, Admixture, |
| I5239 | Balkans_HG | Serbia | Palaeolithic | 10333 | F | 44.60 | 22.01 | Mathieson et al. (2018) doi: 10.1038/nature25778 | PCA, Admixture, |
| I5240 | Balkans_HG | Serbia | Palaeolithic | 10805 | M | 44.60 | 22.01 | Mathieson et al. (2018) doi: 10.1038/nature25778 | PCA, Admixture, |
| I5241 | Balkans_HG | Serbia | Palaeolithic | 11196 | F | 44.60 | 22.01 | Mathieson et al. (2018) doi: 10.1038/nature25778 | PCA, Admixture, |
| I5242 | Balkans_HG | Serbia | Palaeolithic | 10530 | F | 44.60 | 22.01 | Mathieson et al. (2018) doi: 10.1038/nature25778 | PCA, Admixture, NNLS, |
| I5244 | Balkans_HG | Serbia | Palaeolithic | 10785 | F | 44.60 | 22.01 | Mathieson et al. (2018) doi: 10.1038/nature25778 | PCA, Admixture, |
| I5427 | SEEurope_Peloponnese_N | Greece | Neolithic | 7892 | F | 36.64 | 22.38 | Mathieson et al. (2018) doi: 10.1038/nature25778 | PCA, Admixture, |
| I5773_published | Balkans_HG | Serbia | Palaeolithic | 10040 | M | 44.53 | 22.05 | Mathieson et al. (2018) doi: 10.1038/nature25778 | PCA, Admixture, |
| I6601 | SWEurope_C | Portugal | Copper Age | 4250 | M | 39.09 | -9.29 | Olalde et al. (2018) doi: 10.1038/nature25738 | PCA, Admixture, |
| I9005 | SEEurope_Minoan_BA | Greece | Bronze Age | 4000 | F | 35.08 | 25.83 | Lazaridis et al. (2017) doi: 10.1038/nature23310 | PCA, Admixture, |
| I9006 | SEEurope_Mycenaean_BA | Greece | Bronze Age | 3287 | F | 37.97 | 23.50 | Lazaridis et al. (2017) doi: 10.1038/nature23310 | PCA, Admixture, |
| I9010 | SEEurope_Mycenaean_BA | Greece | Bronze Age | 3250 | F | 37.50 | 23.45 | Lazaridis et al. (2017) doi: 10.1038/nature23310 | PCA, Admixture, |

| | | | | | | | | | |
|---|---|---|---|---|---|---|---|---|---|
| I9028.SG | SAfrica_HG | South Africa | Palaeolithic | 2103 | M | -32.76 | 18.03 | Skoglund et al. (2017) doi: 10.1016/j.cell.2017.08.049 | PCA, Admixture, |
| I9033 | SEEurope_Mycenaean_BA | Greece | Bronze Age | 3298 | F | 36.92 | 21.70 | Lazaridis et al. (2017) doi: 10.1038/nature23310 | PCA, Admixture, |
| I9041 | SEEurope_Mycenaean_BA | Greece | Bronze Age | 3250 | M | 37.50 | 23.45 | Lazaridis et al. (2017) doi: 10.1038/nature23310 | PCA, Admixture, |
| I9133.SG | SAfrica_LSA | South Africa | Palaeolithic | 1970 | M | -31.98 | 18.52 | Skoglund et al. (2017) doi: 10.1016/j.cell.2017.08.049 | PCA, Admixture, |
| I9134.SG | SAfrica_LSA | South Africa | Palaeolithic | 1199 | F | -32.81 | 17.95 | Skoglund et al. (2017) doi: 10.1016/j.cell.2017.08.049 | PCA, Admixture, |
| IAM.4 | NAfrica_EN | Morocco | Neolithic | 7021 | M | 33.82 | -6.07 | Fregel et al. (2018) doi: 10.1073/pnas.1800851115 | PCA, Admixture, |
| IAM.5 | NAfrica_EN | Morocco | Neolithic | 7125 | M | 33.82 | -6.07 | Fregel et al. (2018) doi: 10.1073/pnas.1800851115 | PCA, Admixture, Left-pop in qpWave and qpAdm, |
| IAM.6 | NAfrica_EN | Morocco | Neolithic | 6870 | F | 33.82 | -6.07 | Fregel et al. (2018) doi: 10.1073/pnas.1800851115 | PCA, Admixture, |
| IAM.7 | NAfrica_EN | Morocco | Neolithic | 5665 | M | 33.82 | -6.07 | Fregel et al. (2018) doi: 10.1073/pnas.1800851115 | PCA, Admixture, |
| Iboussieres25-1 | WEurope_WHG | France | Palaeolithic | 11725 | M | 44.29 | 4.46 | Mathieson et al. (2018) doi: 10.1038/nature25778 | PCA, Admixture, |
| Iboussieres31-2 | WEurope_WHG | France | Palaeolithic | 11725 | M | 44.29 | 4.46 | Mathieson et al. (2018) doi: 10.1038/nature25778 | PCA, Admixture, |
| Iceman.SG | SEurope_MN | Italy | Neolithic | 5244 | M | 46.77 | 10.83 | Keller et al. (2012) doi: 10.1038/ncomms1701 | PCA, Admixture, NNLS, |
| In661.WGC | SEAsia_LN_BA_IA | Indonesia | Bronze Age | 1872 | F | 0.59 | 101.34 | McColl et al. (2018) doi: 10.1126/science.aat3628 | PCA, Admixture, |
| In662.SG | SEAsia_LN_BA_IA | Indonesia | Bronze Age | 2176 | M | 0.59 | 101.34 | McColl et al. (2018) doi: 10.1126/science.aat3628 | PCA, Admixture, |
| KEB.1 | NAfrica_LN | Morocco | Neolithic | 5650 | F | 34.03 | -6.83 | Fregel et al. (2018) doi: 10.1073/pnas.1800851115 | PCA, Admixture, |
| KEB.4 | NAfrica_LN | Morocco | Neolithic | 5650 | F | 34.03 | -6.83 | Fregel et al. (2018) doi: 10.1073/pnas.1800851115 | PCA, Admixture, Left-pop in qpWave and qpAdm, |
| KEB.6 | NAfrica_LN | Morocco | Neolithic | 5650 | M | 34.03 | -6.83 | Fregel et al. (2018) doi: 10.1073/pnas.1800851115 | PCA, Admixture, |
| KK1.SG | Kotias_HG/Caucasus_HG | Georgia | Palaeolithic | 9720 | M | 42.28 | 43.28 | Jones et al. (2015) doi: 10.1038/ncomms9912 | PCA, Admixture, NNLS, Left-pop in qpWave and qpAdm, Left-pop in qpWave and qpAdm for Basal Eurasia, *f4*-statistics, f4-statistics for Neanderthal |
| Kostenki14 | Russia_EUP | Russia | Palaeolithic | 37470 | M | 51.23 | 39.30 | Fu et al. (2016) doi: 10.1038/nature17993 | PCA, Admixture, Right-pop in qpWave and qpAdm (Outgroup), Right-pop in qpWave and qpAdm for Basal Eurasia (Outgroup), |
| kum6.SG | Anatolia_N | Turkey | Neolithic | 6682 | F | 39.95 | 26.20 | Omrak et al. (2016) doi: 10.1016/j.cub.2015.12.019 | PCA, Admixture, |
| La364.SG | SEAsia_LN_BA | Laos | Bronze Age | 2976 | M | 17.81 | 97.78 | McColl et al. (2018) doi: 10.1126/science.aat3628 | PCA, Admixture, |
| La368.SG | SEAsia_HG | Laos | Palaeolithic | 7872 | M | 18.44 | 104.47 | McColl et al. (2018) doi: 10.1126/science.aat3628 | PCA, Admixture, |
| La727.WGC | SEAsia_BA | Laos | Bronze Age | 2281 | F | 17.81 | 97.78 | McColl et al. (2018) doi: 10.1126/science.aat3628 | PCA, Admixture, |
| Loschbour_published.DG | CEurope_HG | Luxembourg | Palaeolithic | 8050 | M | 49.81 | 6.40 | Lazaridis et al. (2014) doi: 10.1038/nature13673 | PCA, Admixture, NNLS, Right-pop in qpWave and qpAdm (Outgroup), Left-pop in qpWave and qpAdm for Basal Eurasia, *f4*-statistics, *f4*-statistics for Neanderthal |
| LugarCanto41.SG | SWEurope_N | Portugal | Neolithic | 5950 | F | 40.78 | -7.52 | Martiniano et al. (2017) doi: 10.1371/journal.pgen.1006852 | PCA, Admixture, |
| LugarCanto42.SG | SWEurope_N | Portugal | Neolithic | 5950 | F | 40.78 | -7.52 | Martiniano et al. (2017) doi: 10.1371/journal.pgen.1006852 | PCA, Admixture, NNLS, |

| | | | | | | | | | |
|---|---|---|---|---|---|---|---|---|---|
| LugarCanto44.SG | SWEurope_N | Portugal | Neolithic | 5950 | F | 40.78 | -7.52 | Martiniano et al. (2017) doi: 10.1371/journal.pgen.1006852 | PCA, Admixture, |
| LugarCanto45.SG | SWEurope_N | Portugal | Neolithic | 5950 | M | 40.78 | -7.52 | Martiniano et al. (2017) doi: 10.1371/journal.pgen.1006852 | PCA, Admixture, |
| M63.SG | SAsia_Mebrak | Nepal | Historical or Others | 2125 | M | 29.00 | 83.85 | Jeong et al. (2016) doi: 10.1073/pnas.1520844113 | Right-pop in qpWave and qpAdm (Outgroup), |
| MA1.SG | Russia_HG | Russia | Palaeolithic | 24305 | M | 52.90 | 103.50 | Raghavan et al. (2013) doi: 10.1038/nature12736 | PCA, Admixture, Right-pop in qpWave and qpAdm (Outgroup), |
| Ma555.SG | SEAsia_Historical | Malaysia | Historical or Others | 376 | M | 5.25 | 117.00 | McColl et al. (2018) doi: 10.1126/science.aat3628 | Right-pop in qpWave and qpAdm (Outgroup), |
| Ma912.SG | SEAsia_LN | Malaysia | Neolithic | 2520 | M | 5.25 | 102.00 | McColl et al. (2018) doi: 10.1126/science.aat3628 | PCA, Admixture, Right-pop in qpWave and qpAdm (Outgroup), |
| mfo001.SG | SAfrica_IA | South Africa | Iron Age | 378 | F | 28.71 | 30.84 | Schlebusch et al. (2017) doi: 10.1126/science.aao6266 | PCA, Admixture, |
| MonteCanelas337A.SG | SWEurope_N | Portugal | Neolithic | 4953 | M | 37.21 | -8.55 | Martiniano et al. (2017) doi: 10.1371/journal.pgen.1006852 | PCA, Admixture, |
| mota.SG | Mota_EN/EAfrica_EN | Ethiopia | Neolithic | 4472 | M | 6.80 | 38.21 | Llorente et al. (2015) doi: 10.1126/science.aaf3945 | PCA, Admixture, Right-pop in qpWave and qpAdm (Outgroup), Left-pop in qpWave and qpAdm for Basal Eurasia, |
| mur.SG | SWEurope_N | Spain | Neolithic | 7136 | M | 37.54 | -4.30 | Valdiosera et al. (2018) doi: 10.1073/pnas.1717762115 | PCA, Admixture, |
| new001.SG | SAfrica_IA | South Africa | Iron Age | 418 | F | 27.71 | 30.00 | Schlebusch et al. (2017) doi: 10.1126/science.aao6266 | PCA, Admixture, |
| Ostuni1_d | SEurope_MUP | Italy | Palaeolithic | 27620 | F | 40.73 | 17.57 | Fu et al. (2016) doi: 10.1038/nature17993 | PCA, Admixture, |
| Paglicci133_published | SEurope_MUP | Italy | Palaeolithic | 32895 | M | 41.65 | 15.61 | Fu et al. (2016) doi: 10.1038/nature17993 | PCA, Admixture, |
| RISE487.SG | SEurope_BA | Italy | Bronze Age | 5245 | M | 45.26 | 10.38 | Allentoft et al. (2015) doi: 10.1038/nature14507 | PCA, Admixture, |
| RISE489.SG | SEurope_BA | Italy | Bronze Age | 4693 | M | 45.26 | 10.38 | Allentoft et al. (2015) doi: 10.1038/nature14507 | PCA, Admixture, |
| RISE500.SG | RussiaSteppe_MLBA | Russia | Bronze Age | 3550 | F | 53.46 | 85.45 | Allentoft et al. (2015) doi: 10.1038/nature14507 | PCA, Admixture, |
| RISE505.SG | RussiaSteppe_MLBA | Russia | Bronze Age | 3636 | F | 53.46 | 85.45 | Allentoft et al. (2015) doi: 10.1038/nature14507 | PCA, Admixture, NNLS, |
| RISE552.SG | RussiaSteppe_EMBA | Russia | Bronze Age | 4446 | M | 46.62 | 43.33 | Allentoft et al. (2015) doi: 10.1038/nature14507 | PCA, Admixture, |
| S10.SG | SAsia_Samdzong | Nepal | Historical or Others | 1500 | M | 28.93 | 83.91 | Jeong et al. (2016) doi: 10.1073/pnas.1520844113 | Right-pop in qpWave and qpAdm (Outgroup), |
| S35.SG | SAsia_Samdzong | Nepal | Historical or Others | 1500 | M | 28.93 | 83.91 | Jeong et al. (2016) doi: 10.1073/pnas.1520844113 | Right-pop in qpWave and qpAdm (Outgroup), |
| S41.SG | SAsia_Samdzong | Nepal | Historical or Others | 1500 | M | 28.93 | 83.91 | Jeong et al. (2016) doi: 10.1073/pnas.1520844113 | Right-pop in qpWave and qpAdm (Outgroup), |
| san216.SG | SWEurope_N | Spain | Neolithic | 5666 | M | 42.31 | 0.36 | Valdiosera et al. (2018) doi: 10.1073/pnas.1717762115 | PCA, Admixture, |
| SATP.SG | Satsurblia_HG/Caucasus_HG | Georgia | Palaeolithic | 13255 | M | 42.38 | 42.59 | Jones et al. (2015) doi: 10.1038/ncomms9912 | PCA, Admixture, Left-pop in qpWave and qpAdm for Basal Eurasia, *f4*-statistics, , *f4*-statistics for Neanderthal |
| Stuttgart_published.DG | CEurope_LBK_EN | Germany | Neolithic | 7140 | F | 48.78 | 9.18 | Lazaridis et al. (2014) doi: 10.1038/nature13673 | PCA, Admixture, Right-pop in qpWave and qpAdm (Outgroup) |
| TAF009 | Iberomaurusian | Morocco | Palaeolithic | 14357 | M | 34.81 | -2.41 | van de Loosdrecht et al. (2018) doi: 10.1126/science.aar8380 | PCA, Admixture |
| TAF010 | Iberomaurusian | Morocco | Palaeolithic | 14605 | M | 34.81 | -2.41 | van de Loosdrecht et al. (2018) doi: 10.1126/science.aar8380 | PCA, Admixture |
| TAF011 | Iberomaurusian | Morocco | Palaeolithic | 14500 | M | 34.81 | -2.41 | van de Loosdrecht et al. (2018) doi: 10.1126/science.aar8380 | PCA, Admixture, NNLS, Left-pop in qpWave and qpAdm, Left-pop in qpWave and qpAdm for Basal |

| | | | | | | | | | |
|---|---|---|---|---|---|---|---|---|---|
| | | | | | | | | | Eurasia, f4-Statistics, f4-Statistics for Neanderthal |
| TAF012 | Iberomaurusian | Morocco | Palaeolithic | 14470 | F | 34.81 | -2.41 | van de Loosdrecht et al. (2018) doi: 10.1126/science.aar8380 | PCA, Admixture, |
| TAF013 | Iberomaurusian | Morocco | Palaeolithic | 14500 | M | 34.81 | -2.41 | van de Loosdrecht et al. (2018) doi: 10.1126/science.aar8380 | PCA, Admixture, |
| TAF014 | Iberomaurusian | Morocco | Palaeolithic | 14500 | M | 34.81 | -2.41 | van de Loosdrecht et al. (2018) doi: 10.1126/science.aar8380 | PCA, Admixture, |
| TAF015 | Iberomaurusian | Morocco | Palaeolithic | 14500 | M | 34.81 | -2.41 | van de Loosdrecht et al. (2018) doi: 10.1126/science.aar8380 | PCA, Admixture, |
| Tep002.SG | Anatolia_N | Turkey | Neolithic | 8585 | F | 38.17 | 34.49 | Kilinc et al. (2016) doi: 10.1016/j.cub.2016.07.057 | PCA, Admixture, |
| Tep003.SG | Anatolia_N | Turkey | Neolithic | 8505 | M | 38.17 | 34.49 | Kilinc et al. (2016) doi: 10.1016/j.cub.2016.07.057 | PCA, Admixture, |
| Tep004.SG | Anatolia_N | Turkey | Neolithic | 8295 | F | 38.17 | 34.49 | Kilinc et al. (2016) doi: 10.1016/j.cub.2016.07.057 | PCA, Admixture, |
| Tep006.SG | Anatolia_N | Turkey | Neolithic | 8230 | M | 38.17 | 34.49 | Kilinc et al. (2016) doi: 10.1016/j.cub.2016.07.057 | PCA, Admixture, |
| Tianyuan | EAsia_UP | China | Palaeolithic | 39475 | M | 39.39 | 115.52 | Yang et al. (2017) doi: 10.1016/j.cub.2017.09.030 | PCA, Admixture, |
| TOR.11 | SWEurope_N | Spain | Neolithic | 6980 | F | 36.63 | -4.52 | Fregel et al. (2018) doi: 10.1073/pnas.1800851115 | PCA, Admixture, |
| TOR.6 | SWEurope_N | Spain | Neolithic | 6980 | F | 36.63 | -4.52 | Fregel et al. (2018) doi: 10.1073/pnas.1800851115 | PCA, Admixture, |
| TOR.7 | SWEurope_N | Spain | Neolithic | 6980 | F | 36.63 | -4.52 | Fregel et al. (2018) doi: 10.1073/pnas.1800851115 | PCA, Admixture, |
| TOR.8 | SWEurope_N | Spain | Neolithic | 6980 | F | 36.63 | -4.52 | Fregel et al. (2018) doi: 10.1073/pnas.1800851115 | PCA, Admixture, |
| Ust_Ishim_published.DG | Ust_Ishim/Russia_HG | Russia | Palaeolithic | 45020 | M | 57.70 | 71.10 | Fu et al. (2014) doi: 10.1038/nature13810 | PCA, Admixture, Right-pop in qpWave and qpAdm (Outgroup), Right-pop in qpWave and qpAdm for Basal Eurasia (Outgroup), f4-Statistics |
| Vestonice16 | CEurope_MUP | Czech Republic | Palaeolithic | 30010 | M | 48.53 | 16.39 | Fu et al. (2016) doi: 10.1038/nature17993 | PCA, Admixture, Right-pop in qpWave and qpAdm (Outgroup) |
| Villabruna | SEurope_HG | Italy | Palaeolithic | 13980 | M | 46.15 | 12.21 | Fu et al. (2016) doi: 10.1038/nature17993 | PCA, Admixture, NNLS |
| Vt777.SG | SEAsia_N | Vietnam | Neolithic | 2265 | F | 19.81 | 105.79 | McColl et al. (2018) doi: 10.1126/science.aat3628 | PCA, Admixture |
| Vt778.SG | SEAsia_N | Vietnam | Neolithic | 2625 | M | 22.39 | 103.47 | McColl et al. (2018) doi: 10.1126/science.aat3628 | PCA, Admixture |
| Vt779.SG | SEAsia_BA | Vietnam | Bronze Age | 2247 | F | 19.81 | 105.79 | McColl et al. (2018) doi: 10.1126/science.aat3628 | PCA, Admixture |
| Vt781.SG | SEAsia_BA | Vietnam | Bronze Age | 2249 | F | 19.81 | 105.79 | McColl et al. (2018) doi: 10.1126/science.aat3628 | PCA, Admixture |
| Vt796.SG | SEAsia_BA | Vietnam | Bronze Age | 2172 | F | 19.81 | 105.79 | McColl et al. (2018) doi: 10.1126/science.aat3628 | PCA, Admixture |
| Vt808.SG | SEAsia_BA | Vietnam | Bronze Age | 2251 | F | 19.81 | 105.79 | McColl et al. (2018) doi: 10.1126/science.aat3628 | PCA, Admixture |
| Vt833.SG | SEAsia_N | Vietnam | Neolithic | 4149 | F | 19.81 | 105.79 | McColl et al. (2018) doi: 10.1126/science.aat3628 | PCA, Admixture |
| Vt880.SG | SEAsia_N | Vietnam | Neolithic | 4000 | F | 21.01 | 107.29 | McColl et al. (2018) doi: 10.1126/science.aat3628 | PCA, Admixture |
| WC1.SG | Zagros_N | Iran | Neolithic | 9219 | M | 34.61 | 47.11 | Broushaki et al. (2016) doi: 10.1126/science.aaf7943 | PCA, Admixture, NNLS, Left-pop in qpWave and qpAdm, Left-pop in qpWave and qpAdm for Basal Eurasia, f4-Statistics, f4-Statistics for Neanderthal |

**Table S3:** ‘Right’ populations used qpAdm for all the possible source combinations of N= (2..5)

| SampleID | Group ID | Publication |
|---|---|---|
| Bichon.SG | CEurope_HG | Jones et al. (2015) doi: 10.1038/ncomms9912 |
| DA38.SG | CAsia_XiongNu_WE | Damgaard et al. (2018) doi:10.1038/s41586-018-0094-2 |
| DA39.SG | CAsia_XiongNu | Damgaard et al. (2018) doi:10.1038/s41586-018-0094-2 |
| DA43.SG | CAsia_XiongNu | Damgaard et al. (2018) doi:10.1038/s41586-018-0094-2 |
| DA45.SG | CAsia_XiongNu | Damgaard et al. (2018) doi:10.1038/s41586-018-0094-2 |
| I0100 | CEurope_LBK_EN | Lipson et al. (2017) doi: 10.1038/nature24476 |
| I0103 | CEurope_LN | Mathieson et al. (2015) doi: 10.1038/nature16152 |
| I0112 | CEurope_BellBeaker_BA | Mathieson et al. (2015) doi: 10.1038/nature16152 |
| I1499 | CEurope_MN | Mathieson et al. (2015) doi: 10.1038/nature16152 |
| Kostenki14 | Russia_EUP | Fu et al. (2016) doi: 10.1038/nature17993 |
| Loschbour_published.DG | Loschbour_HG/CEurope_HG | Lazaridis et al. (2014) doi: 10.1038/nature13673 |
| M63.SG | SAsia_Mebrak | Jeong et al. (2016) doi: 10.1073/pnas.1520844113 |
| MA1.SG | Russia_HG | Raghavan et al. (2013) doi: 10.1038/nature12736 |
| Ma555.SG | SEAsia_Historical | McColl et al. (2018) doi: 10.1126/science.aat3628 |
| Ma912.SG | SEAsia_LN | McColl et al. (2018) doi: 10.1126/science.aat3628 |
| mota.SG | Mota_EN/EAfrica_EN | Llorente et al. (2015) doi: 10.1126/science.aaf3945 |
| S10.SG | SAsia_Samdzong | Jeong et al. (2016) doi: 10.1073/pnas.1520844113 |
| S35.SG | SAsia_Samdzong | Jeong et al. (2016) doi: 10.1073/pnas.1520844113 |
| S41.SG | SAsia_Samdzong | Jeong et al. (2016) doi: 10.1073/pnas.1520844113 |
| Stuttgart_published.DG | CEurope_LBK_EN | Lazaridis et al. (2014) doi: 10.1038/nature13673 |
| Ust_Ishim_published.DG | Ust_Ishim/Russia_HG | Fu et al. (2014) doi: 10.1038/nature13810 |
| Vestonice16 | CEurope_MUP | Fu et al. (2016) doi: 10.1038/nature17993 |

**Table S4:** Right populations used qpAdm to search for traces of the basal Eurasian lineage in modern AP populations in the form (Test, Mota, Early hunter-gatherer)

| SampleID | Group ID | Publication |
|---|---|---|
| Kostenki14 | Russia_EUP | Fu et al. (2016) doi: 10.1038/nature17993 |
| Ust_Ishim_published.DG | Ust_Ishim/Russia_HG | Fu et al. (2014) doi: 10.1038/nature13810 |
| BR_Onge-2.DG | Onge | Mallick et al. (2016) doi: 10.1038/nature18964 |
| S_Han-1.DG | Han | Mallick et al. (2016) doi: 10.1038/nature18964 |
| S_Papuan-14.DG | Papuan | Skoglund et al. (2015) doi: 10.1038/nature14895 |

**Table S5:** Mixture proportions on AP and Iran populations inferred by *qpAdm* as a combination of four ancient sources

| Target | Pop1 | Pop2 | Pop3 | Pop4 | dof | chisq | tail | Prob.Pop1 | Prob.Pop2 | Prob.Pop3 | Prob.Pop4 | Feasible | SE.Pop1 | SE.Pop2 | SE.Pop3 | SE.Pop4 |
|---|---|---|---|---|---|---|---|---|---|---|---|---|---|---|---|---|
| M_Yemen | Caucasus_HG | Zagros_N | Iberomaurusian | Natufian | 7 | 10.36 | 0.16888 | 0.105 | 0.105 | 0.12 | 0.67 | Yes | 0.85 | 0.664 | 0.284 | 0.475 |
| M_Yemen | Caucasus_HG | Levant_EBA | Levant_PPN | Iberomaurusian | 7 | 5.748 | 0.56951 | 0.06 | 0.625 | 0.03 | 0.285 | Yes | 0.205 | 0.411 | 0.241 | 0.038 |
| M_Iran | Caucasus_EBA | Caucasus_HG | Zagros_N | Iberomaurusian | 7 | 11.31 | 0.12577 | 0.413 | 0.286 | 0.179 | 0.123 | Yes | 1.038 | 1.621 | 0.657 | 0.076 |
| M_Iran | Caucasus_HG | Zagros_N | Levant_PPN | Iberomaurusian | 7 | 3.126 | 0.87308 | 0.458 | 0.358 | 0.157 | 0.027 | Yes | 0.215 | 0.216 | 0.062 | 0.06 |
| M_Iran | Caucasus_HG | Zagros_N | Levant_EBA | Iberomaurusian | 7 | 4.858 | 0.67735 | 0.593 | 0.164 | 0.151 | 0.092 | Yes | 0.17 | 0.162 | 0.08 | 0.051 |
| M_Iran | Caucasus_HG | Zagros_N | Iberomaurusian | Natufian | 7 | 5.025 | 0.65685 | 0.519 | 0.246 | 0.039 | 0.196 | Yes | 0.238 | 0.209 | 0.091 | 0.135 |
| M_UAE | Caucasus_EBA | Caucasus_HG | Zagros_N | Iberomaurusian | 7 | 9.024 | 0.25093 | 0.393 | 0.104 | 0.217 | 0.287 | Yes | 0.312 | 0.497 | 0.232 | 0.049 |
| M_UAE | Caucasus_HG | Zagros_N | Levant_EBA | Iberomaurusian | 7 | 5.567 | 0.59109 | 0.437 | 0.173 | 0.141 | 0.249 | Yes | 0.158 | 0.153 | 0.07 | 0.047 |
| M_UAE | Caucasus_HG | Zagros_N | Levant_PPN | Iberomaurusian | 7 | 4.513 | 0.71914 | 0.329 | 0.344 | 0.13 | 0.198 | Yes | 0.236 | 0.233 | 0.065 | 0.061 |
| M_UAE | Caucasus_HG | Zagros_N | Iberomaurusian | Natufian | 7 | 3.593 | 0.82531 | 0.472 | 0.175 | 0.235 | 0.118 | Yes | 0.168 | 0.151 | 0.067 | 0.101 |
| M_Oman | Caucasus_HG | Zagros_N | Levant_PPN | Iberomaurusian | 7 | 5.712 | 0.57374 | 0.181 | 0.417 | 0.214 | 0.188 | Yes | 0.251 | 0.253 | 0.065 | 0.066 |
| M_Oman | Caucasus_HG | Zagros_N | Levant_EBA | Iberomaurusian | 7 | 7.293 | 0.39898 | 0.296 | 0.171 | 0.265 | 0.268 | Yes | 0.177 | 0.167 | 0.072 | 0.049 |
| M_Oman | Caucasus_HG | Zagros_N | Iberomaurusian | Natufian | 7 | 5.556 | 0.59242 | 0.371 | 0.198 | 0.215 | 0.216 | Yes | 0.208 | 0.181 | 0.077 | 0.118 |
| M_SaudiArabia | Caucasus_EBA | Zagros_N | Levant_PPN | Iberomaurusian | 7 | 4.271 | 0.74807 | 0.359 | 0.118 | 0.34 | 0.183 | Yes | 0.394 | 0.333 | 0.19 | 0.128 |

| | | | | | | | | | | | | | | | | |
|---|---|---|---|---|---|---|---|---|---|---|---|---|---|---|---|---|
| M_SaudiArabia | Caucasus_HG | Zagros_N | Levant_PPN | Iberomaurusian | 7 | 4.441 | 0.7278 | 0.1 | 0.35 | 0.468 | 0.082 | Yes | 0.223 | 0.229 | 0.062 | 0.066 |
| M_SaudiArabia | Caucasus_HG | Zagros_N | Iberomaurusian | Natufian | 7 | 6.984 | 0.43059 | 0.169 | 0.127 | 0.034 | 0.67 | Yes | 0.306 | 0.251 | 0.111 | 0.178 |
| M_SaudiArabia | Caucasus_HG | Levant_EBA | Levant_PPN | Iberomaurusian | 7 | 5.335 | 0.6191 | 0.229 | 0.451 | 0.129 | 0.191 | Yes | 0.163 | 0.33 | 0.197 | 0.032 |
| M_SaudiArabia | Zagros_N | Levant_EBA | Levant_PPN | Iberomaurusian | 7 | 5.828 | 0.56002 | 0.268 | 0.338 | 0.27 | 0.123 | Yes | 7.816 | 14.98 | 9.988 | 2.825 |
| M_SaudiArabia | Zagros_N | Levant_EBA | Levant_PPN | Natufian | 7 | 3.211 | 0.86482 | 0.267 | 0.072 | 0.132 | 0.53 | Yes | 0.106 | 0.187 | 0.267 | 0.321 |

**Table S6:** Mixture proportions on AP and Iran populations inferred by *qpAdm* for all the possible source combinations of N= (2..5)

| Pop2 | dof | chisq | tail | Prob.Pop1 | Prob.Pop2 | Feasible | SE.Pop1 | SE.Pop2 |
|---|---|---|---|---|---|---|---|---|
| Natufian | 9 | 21.657 | 0.0100 | 0.032 | 0.97 | Yes | 0.274 | 0.274 |
| Levant_PPN | 9 | 21.504 | 0.0106 | 0.552 | 0.45 | Yes | 0.04 | 0.04 |
| Natufian | 9 | 12.543 | 0.1844 | 0.254 | 0.75 | Yes | 0.073 | 0.073 |
| Iberomaurusian | 9 | 6.738 | 0.6644 | 0.708 | 0.29 | Yes | 0.031 | 0.031 |
| Natufian | 9 | 14.253 | 0.1136 | 0.155 | 0.85 | Yes | 0.055 | 0.055 |
| Zagros_N | 9 | 15.734 | 0.0727 | 0.474 | 0.53 | Yes | 0.077 | 0.077 |
| Iberomaurusian | 9 | 15.444 | 0.0794 | 0.779 | 0.22 | Yes | 0.035 | 0.035 |
| Zagros_N | 9 | 12.248 | 0.1997 | 0.728 | 0.27 | Yes | 0.134 | 0.134 |
| Levant_PPN | 9 | 12.864 | 0.1689 | 0.971 | 0.03 | Yes | 0.074 | 0.074 |
| Iberomaurusian | 9 | 9.135 | 0.4249 | 0.888 | 0.11 | Yes | 0.037 | 0.037 |
| Natufian | 9 | 15.121 | 0.0877 | 0.888 | 0.11 | Yes | 0.171 | 0.171 |
| Levant_EBA | 9 | 18.097 | 0.0341 | 0.614 | 0.39 | Yes | 0.074 | 0.074 |
| Levant_PPN | 9 | 9.635 | 0.3808 | 0.722 | 0.28 | Yes | 0.04 | 0.04 |
| Natufian | 9 | 18.21 | 0.0328 | 0.556 | 0.44 | Yes | 0.097 | 0.097 |
| Iberomaurusian | 9 | 13.744 | 0.1317 | 0.619 | 0.38 | Yes | 0.032 | 0.032 |
| Iberomaurusian | 9 | 10.085 | 0.3437 | 0.72 | 0.28 | Yes | 0.034 | 0.034 |
| Levant_PPN | 9 | 13.526 | 0.1402 | 0.808 | 0.19 | Yes | 0.044 | 0.044 |
| Natufian | 9 | 14.927 | 0.0930 | 0.624 | 0.38 | Yes | 0.091 | 0.091 |
| Iberomaurusian | 9 | 9.171 | 0.4216 | 0.622 | 0.38 | Yes | 0.031 | 0.031 |
| Iberomaurusian | 9 | 16.861 | 0.0509 | 0.718 | 0.28 | Yes | 0.035 | 0.035 |
| Levant_PPN | 9 | 15.014 | 0.0905 | 0.768 | 0.23 | Yes | 0.044 | 0.044 |
| Natufian | 9 | 13.943 | 0.1244 | 0.571 | 0.43 | Yes | 0.087 | 0.087 |
| Iberomaurusian | 9 | 18.341 | 0.0314 | 0.793 | 0.21 | Yes | 0.035 | 0.035 |
| Natufian | 9 | 14.019 | 0.1217 | 0.077 | 0.92 | Yes | 0.242 | 0.242 |
| Natufian | 9 | 11.711 | 0.2301 | 0.271 | 0.73 | Yes | 0.141 | 0.141 |
| Levant_EBA | 9 | 18.947 | 0.0256 | 0.323 | 0.68 | Yes | 0.066 | 0.066 |

| | | | | | | | | |
|---|---|---|---|---|---|---|---|---|
| Levant_PPN | 9 | 6.158 | 0.7240 | 0.513 | 0.49 | Yes | 0.036 | 0.036 |
| Natufian | 9 | 7.382 | 0.5975 | 0.239 | 0.76 | Yes | 0.072 | 0.072 |
| Iberomaurusian | 9 | 7.874 | 0.5469 | 0.781 | 0.22 | Yes | 0.032 | 0.032 |
| Natufian | 9 | 13.769 | 0.1308 | 0.069 | 0.93 | Yes | 0.059 | 0.059 |

| **Pop2** | **Pop3** | **dof** | **chisq** | **tail** | **Prob.Pop1** | **Prob.Pop2** | **Prob.Pop3** | **Feasible** | **SE.Pop1** | **SE.Pop2** | **SE.Pop3** |
|---|---|---|---|---|---|---|---|---|---|---|---|
| Caucasus_HG | Iberomaurusian | 8 | 12.454 | 0.1321 | 0.22 | 0.459 | 0.324 | Yes | 1.384 | 1.54 | 0.2 |
| Zagros_N | Iberomaurusian | 8 | 9.055 | 0.3377 | 0.46 | 0.257 | 0.285 | Yes | 0.078 | 0.12 | 0.1 |
| Zagros_N | Iberomaurusian | 8 | 9.542 | #### | 0.64 | 0.105 | 0.253 | Yes | 0.137 | 0.18 | 0.1 |
| Levant_EBA | Iberomaurusian | 8 | 7.149 | 0.5207 | 0.6 | 0.11 | 0.292 | Yes | 0.096 | 0.08 | 0 |
| Levant_PPN | Iberomaurusian | 8 | 7.288 | 0.5059 | 0.67 | 0.053 | 0.281 | Yes | 0.063 | 0.05 | 0 |
| Iberomaurusian | Natufian | 8 | 5.144 | 0.7421 | 0.65 | 0.3 | 0.05 | Yes | 0.099 | 0.04 | 0.1 |
| Levant_EBA | Iberomaurusian | 8 | 14.763 | #### | 0.54 | 0.295 | 0.169 | Yes | 0.111 | 0.07 | 0.1 |
| Levant_PPN | Iberomaurusian | 8 | 7.804 | 0.4529 | 0.66 | 0.21 | 0.129 | Yes | 0.065 | 0.04 | 0 |
| Iberomaurusian | Natufian | 8 | 12.098 | 0.1469 | 0.57 | 0.079 | 0.356 | Yes | 0.093 | 0.05 | 0.1 |
| Caucasus_HG | Iberomaurusian | 8 | 12.914 | 0.1149 | 0.22 | 0.631 | 0.152 | Yes | 0.464 | 0.51 | 0.1 |
| Zagros_N | Levant_PPN | 8 | 8.062 | 0.4274 | 0.2 | 0.609 | 0.191 | Yes | 0.176 | 0.09 | 0.1 |
| Zagros_N | Iberomaurusian | 8 | 11.333 | 0.1835 | 0.59 | 0.293 | 0.115 | Yes | 0.099 | 0.15 | 0.1 |
| Zagros_N | Natufian | 8 | 15.027 | 0.0586 | 0.1 | 0.497 | 0.404 | Yes | 0.279 | 0.13 | 0.2 |
| Zagros_N | Levant_EBA | 8 | 7.693 | #### | 0.5 | 0.356 | 0.147 | Yes | 0.165 | 0.1 | 0.1 |
| Zagros_N | Levant_PPN | 8 | 2.598 | 0.9570 | 0.39 | 0.442 | 0.173 | Yes | 0.134 | 0.1 | 0 |
| Zagros_N | Iberomaurusian | 8 | 8.981 | #### | 0.83 | 0.075 | 0.093 | Yes | 0.15 | 0.2 | 0.1 |
| Zagros_N | Natufian | 8 | 5.183 | 0.7379 | 0.44 | 0.322 | 0.234 | Yes | 0.126 | 0.09 | 0.1 |
| Levant_EBA | Iberomaurusian | 8 | 5.884 | #### | 0.75 | 0.12 | 0.132 | Yes | 0.104 | 0.09 | 0 |
| Levant_PPN | Iberomaurusian | 8 | 6.138 | 0.6318 | 0.81 | 0.08 | 0.114 | Yes | 0.07 | 0.06 | 0 |
| Iberomaurusian | Natufian | 8 | 7.216 | 0.5135 | 0.78 | 0.133 | 0.088 | Yes | 0.117 | 0.05 | 0.1 |
| Zagros_N | Iberomaurusian | 8 | 8.337 | 0.4013 | 0.55 | 0.11 | 0.336 | Yes | 0.081 | 0.12 | 0.1 |
| Levant_EBA | Iberomaurusian | 8 | 9.004 | #### | 0.46 | 0.234 | 0.311 | Yes | 0.091 | 0.08 | 0 |
| Levant_PPN | Iberomaurusian | 8 | 9.841 | 0.2764 | 0.59 | 0.12 | 0.289 | Yes | 0.061 | 0.05 | 0 |
| Iberomaurusian | Natufian | 8 | 7.456 | #### | 0.58 | 0.29 | 0.131 | Yes | 0.104 | 0.04 | 0.1 |

| | | | | | | | | | | | | | | | | | |
|---|---|---|---|---|---|---|---|---|---|---|---|---|---|---|---|---|---|
| Levant_EBA | Iberomaurusian | 8 | 11.024 | #### | 0.41 | 0.374 | 0.218 | Yes | 0.087 | 0.06 | 0 | | | | | | |
| Levant_PPN | Iberomaurusian | 8 | 6.568 | 0.5839 | 0.59 | 0.257 | 0.151 | Yes | 0.061 | 0.04 | 0 | | | | | | |
| Iberomaurusian | Natufian | 8 | 10.142 | 0.2552 | 0.5 | 0.097 | 0.408 | Yes | 0.088 | 0.05 | 0.1 | | | | | | |
| Levant_EBA | Iberomaurusian | 8 | 6.486 | 0.5929 | 0.2 | 0.594 | 0.206 | Yes | 0.275 | 0.28 | 0 | | | | | | |
| Levant_PPN | Iberomaurusian | 8 | 4.669 | 0.7923 | 0.49 | 0.281 | 0.228 | Yes | 0.074 | 0.06 | 0 | | | | | | |
| Iberomaurusian | Natufian | 8 | 10.638 | 0.2231 | 0.29 | 0.112 | 0.594 | Yes | 0.198 | 0.05 | 0.2 | | | | | | |
| Zagros_N | Natufian | 8 | 6.869 | 0.5508 | 0.1 | 0.193 | 0.706 | Yes | 0.155 | 0.1 | 0.1 | | | | | | |
| Levant_EBA | Iberomaurusian | 8 | 4.87 | 0.7714 | 0.19 | 0.62 | 0.191 | Yes | 0.096 | 0.09 | 0 | | | | | | |
| Levant_PPN | Iberomaurusian | 8 | 8.786 | 0.3607 | 0.43 | 0.399 | 0.17 | Yes | 0.053 | 0.04 | 0 | | | | | | |
| Iberomaurusian | Natufian | 8 | 7.681 | 0.4652 | 0.3 | 0.082 | 0.622 | Yes | 0.107 | 0.04 | 0.1 | | | | | | |
| Levant_EBA | Iberomaurusian | 8 | 6.931 | 0.5441 | 0.09 | 0.728 | 0.179 | Yes | 0.096 | 0.06 | 0.1 | | | | | | |
| Levant_EBA | Natufian | 8 | 4.067 | 0.8510 | 0.23 | 0.014 | 0.756 | Yes | 0.067 | 0.25 | 0.3 | | | | | | |
| Levant_PPN | Iberomaurusian | 8 | 4.615 | 0.7978 | 0.45 | 0.494 | 0.06 | Yes | 0.059 | 0.04 | 0 | | | | | | |
| Levant_PPN | Natufian | 8 | 5.212 | 0.7347 | 0.35 | 0.298 | 0.348 | Yes | 0.205 | 0.52 | 0.7 | | | | | | |
| Levant_PPN | Iberomaurusian | 8 | 9.129 | 0.3315 | 0.4 | 0.299 | 0.3 | Yes | 0.084 | 0.07 | 0 | | | | | | |
| Iberomaurusian | Natufian | 8 | 12.208 | 0.1422 | 0.19 | 0.185 | 0.624 | Yes | 0.215 | 0.05 | 0.2 | | | | | | |
| Levant_EBA | Iberomaurusian | 8 | 6.051 | 0.6415 | 0.09 | 0.632 | 0.279 | Yes | 0.101 | 0.09 | 0 | | | | | | |
| Levant_PPN | Iberomaurusian | 8 | 14.895 | 0.0612 | 0.35 | 0.4 | 0.255 | Yes | 0.056 | 0.04 | 0 | | | | | | |
| Iberomaurusian | Natufian | 8 | 10.653 | 0.2221 | 0.22 | 0.162 | 0.616 | Yes | 0.117 | 0.04 | 0.1 | | | | | | |
| Levant_EBA | Iberomaurusian | 8 | 6.613 | 0.5789 | 0.03 | 0.691 | 0.279 | Yes | 0.098 | 0.06 | 0.1 | | | | | | |
| Levant_PPN | Iberomaurusian | 8 | 9.229 | #### | 0.37 | 0.47 | 0.159 | Yes | 0.06 | 0.03 | 0 | | | | | | |
| Iberomaurusian | Natufian | 8 | 10.365 | #### | 0.19 | 0.086 | 0.729 | Yes | 0.093 | 0.06 | 0.1 | | | | | | |
| **Pop2** | **Pop3** | **Pop4** | **Pop5** | **dof** | **chisq** | **tail** | **Prob.Pop1** | **Prob.Pop2** | **Prob.Pop3** | **Prob.Pop4** | **Prob.Pop5** | **Feasible** | **SE.Pop1** | **SE.Pop2** | **SE.Pop3** | **SE.Pop4** | **SE.Pop5** |
| Zagros_N | Levant_EBA | Levant_PPN | Iberomaurusian | 6 | 3.01 | 0.8074 | 0.531 | 0.236 | 0.107 | 0.06 | 0.1 | Yes | 0.197 | 0.374 | 0.46 | 0.34 | 0.116 |
| Zagros_N | Levant_EBA | Levant_PPN | Iberomaurusian | 6 | 4.76 | 0.5748 | 0.149 | 0.143 | 0.32 | 0.24 | 0.1 | Yes | 0.215 | 0.434 | 0.585 | 0.428 | 0.142 |

**Table S7:** CP/NNLS - Weighted jackknife bootstraps for overall by country – limiting to the four sources identified in the qpAdm results

| | Caucasus_HG | Caucasus_HG-SE | Iberomaurusian | Iberomaurusian-SE | Natufians | Natufians-SE | Zagros_N | Zagros_N-SE |
|---|---|---|---|---|---|---|---|---|
| Iran | 0.49796 | 0.03972 | 0.11231 | 0.01412 | 0.17329 | 0.02679 | 0.21644 | 0.03381 |
| UAE | 0.30593 | 0.02787 | 0.28737 | 0.01379 | 0.12553 | 0.02590 | 0.28117 | 0.02623 |
| Oman | 0.27965 | 0.02317 | 0.26628 | 0.01338 | 0.21558 | 0.02277 | 0.23849 | 0.02582 |
| Saudi Arabia | 0.16309 | 0.02724 | 0.04043 | 0.01533 | 0.64661 | 0.02718 | 0.14988 | 0.04124 |
| Yemen | 0.24566 | 0.03173 | 0.15767 | 0.01330 | 0.52793 | 0.02437 | 0.06874 | 0.04289 |

**Table S8:** CP/NNLS - Weighted jackknife bootstraps for overall by Cluster – limiting to the four sources identified in the qpAdm results

| | Caucasus_HG | Caucasus_HG-SE | Iberomaurusian | Iberomaurusian-SE | Natufians | Natufians-SE | Zagros_N | Zagros_N-SE |
|---|---|---|---|---|---|---|---|---|
| Cluster A | 0.366975412 | 0.03824062 | 0.59111635 | 0.011239602 | 0.0017743 | 0.0107156 | 0.040134 | 0.0383911 |
| Cluster B | 0.586148121 | 0.02661906 | 0 | 0 | 0.4138519 | 0.0266191 | 0 | 0 |
| Cluster C | 0.497906598 | 0.02312623 | 0.073116912 | 0.01555024 | 0.4289765 | 0.0274358 | 0 | 0 |
| Cluster D | 0.615670938 | 0.126787578 | 0.19664222 | 0.018059174 | 0 | 0 | 0.1876868 | 0.1395406 |
| Cluster E | 0.789207075 | 0.024338345 | 0 | 0 | 0.2107929 | 0.0243383 | 0 | 0 |
| Cluster F | 0.181632249 | 0.079475129 | 0 | 0 | 0.7167268 | 0.0238446 | 0.1016409 | 0.0776371 |

**Table S9:** Neanderthal introgression measured through the *f4*(Test, Mbuti; Altai, Denisovan) test

| Pop1_W | Pop2_X | Pop3_Y | Pop4_Z | *f4* statistic | SD | Z |
|---|---|---|---|---|---|---|
| M_Armenia | Mbuti | Altai | Denisova | 0.00107 | 0.000305 | 3.506 |
| M_Bedouin | Mbuti | Altai | Denisova | 0.001042 | 0.000293 | 3.55 |
| M_Druze_Israel | Mbuti | Altai | Denisova | 0.001065 | 0.000298 | 3.567 |
| M_Finland | Mbuti | Altai | Denisova | 0.001253 | 0.000306 | 4.095 |
| M_France | Mbuti | Altai | Denisova | 0.001239 | 0.000308 | 4.021 |
| M_Georgia | Mbuti | Altai | Denisova | 0.001144 | 0.000305 | 3.755 |
| M_Iran | Mbuti | Altai | Denisova | 0.000987 | 0.000293 | 3.367 |
| M_Italy | Mbuti | Altai | Denisova | 0.001222 | 0.000303 | 4.038 |
| M_Jordan | Mbuti | Altai | Denisova | 0.001027 | 0.000296 | 3.469 |
| M_Lebanon | Mbuti | Altai | Denisova | 0.000965 | 0.000308 | 3.134 |
| M_Lezgins | Mbuti | Altai | Denisova | 0.001187 | 0.000305 | 3.896 |
| M_Oman | Mbuti | Altai | Denisova | 0.000902 | 0.000287 | 3.138 |
| M_Romania | Mbuti | Altai | Denisova | 0.001238 | 0.000303 | 4.087 |
| M_SaudiArabia | Mbuti | Altai | Denisova | 0.000968 | 0.000292 | 3.319 |
| M_Spain | Mbuti | Altai | Denisova | 0.001256 | 0.0003 | 4.186 |
| M_Syria | Mbuti | Altai | Denisova | 0.001028 | 0.000301 | 3.41 |
| M_Turkey | Mbuti | Altai | Denisova | 0.001122 | 0.000303 | 3.707 |
| M_UAE | Mbuti | Altai | Denisova | 0.000952 | 0.000287 | 3.315 |
| M_UK | Mbuti | Altai | Denisova | 0.001269 | 0.000307 | 4.131 |
| M_Yemen | Mbuti | Altai | Denisova | 0.000953 | 0.000289 | 3.292 |
| Kotias_HG | Mbuti | Altai | Denisova | 0.000878 | 0.000409 | 2.146 |
| Loschbour | Mbuti | Altai | Denisova | 0.00105 | 0.00038 | 2.765 |
| Iberomaurusian | Mbuti | Altai | Denisova | 0.000645 | 0.000441 | 1.463 |
| Natufian | Mbuti | Altai | Denisova | 0.000798 | 0.000641 | 1.244 |
| Satsurblia_HG | Mbuti | Altai | Denisova | 0.001473 | 0.000469 | 3.142 |
| Zagros_N | Mbuti | Altai | Denisova | 0.000737 | 0.000398 | 1.851 |
| Cluster_A | Mbuti | Altai | Denisova | 0.000775 | 0.000277 | 2.793 |
| Cluster_D | Mbuti | Altai | Denisova | 0.000946 | 0.000295 | 3.206 |
| Cluster_F | Mbuti | Altai | Denisova | 0.001008 | 0.000293 | 3.439 |

**Table S10:** *qpAdm* in the form (Test, Mota, early hunter-gatherer)

| Test | Mota_EN | NWRussia_HG | dof | chisq | tail | Prob. Mota_EN | Prob. NWRussia_HG | Feasible | SE. Mota_EN | SE. NWRussia_HG |
|---|---|---|---|---|---|---|---|---|---|---|
| M_Armenia | Mota_EN | NWRussia_HG | 3 | 5.35 | 0.1479 | 0.26 | 0.74 | Yes | 0.069 | 0.069 |
| M_Bedouin | Mota_EN | NWRussia_HG | 3 | 4.051 | 0.2560 | 0.322 | 0.678 | Yes | 0.063 | 0.063 |
| M_Druze | Mota_EN | NWRussia_HG | 3 | 3.009 | 0.3902 | 0.268 | 0.732 | Yes | 0.067 | 0.067 |
| M_Dubai | Mota_EN | NWRussia_HG | 3 | 5.081 | 0.1660 | 0.447 | 0.553 | Yes | 0.06 | 0.06 |
| M_France | Mota_EN | NWRussia_HG | 3 | 7.066 | 0.0698 | 0.154 | 0.846 | Yes | 0.074 | 0.074 |
| M_Georgia | Mota_EN | NWRussia_HG | 3 | 3.432 | 0.3297 | 0.259 | 0.741 | Yes | 0.066 | 0.066 |
| M_Iran | Mota_EN | NWRussia_HG | 3 | 6.806 | 0.0783 | 0.371 | 0.629 | Yes | 0.066 | 0.066 |
| M_Italy | Mota_EN | NWRussia_HG | 3 | 4.477 | 0.2144 | 0.184 | 0.816 | Yes | 0.07 | 0.07 |
| M_Jordan | Mota_EN | NWRussia_HG | 3 | 4.403 | 0.2211 | 0.334 | 0.666 | Yes | 0.064 | 0.064 |
| M_Lezgins | Mota_EN | NWRussia_HG | 3 | 6.343 | 0.0960 | 0.214 | 0.786 | Yes | 0.071 | 0.071 |
| M_Lebanon | Mota_EN | NWRussia_HG | 3 | 4.62 | 0.2018 | 0.32 | 0.68 | Yes | 0.069 | 0.069 |
| M_Oman | Mota_EN | NWRussia_HG | 3 | 3.003 | 0.3912 | 0.455 | 0.545 | Yes | 0.058 | 0.058 |
| M_Palestinia | Mota_EN | NWRussia_HG | 3 | 2.764 | 0.4295 | 0.332 | 0.668 | Yes | 0.062 | 0.062 |
| M_Romania | Mota_EN | NWRussia_HG | 3 | 3.512 | 0.3192 | 0.203 | 0.797 | Yes | 0.067 | 0.067 |
| M_SaudiArabia | Mota_EN | NWRussia_HG | 3 | 2.641 | 0.4503 | 0.366 | 0.634 | Yes | 0.061 | 0.061 |
| M_Spain | Mota_EN | NWRussia_HG | 3 | 6.649 | 0.0840 | 0.158 | 0.842 | Yes | 0.072 | 0.072 |
| M_Syria | Mota_EN | NWRussia_HG | 3 | 2.963 | 0.3974 | 0.31 | 0.69 | Yes | 0.065 | 0.065 |
| M_UK | Mota_EN | NWRussia_HG | 3 | 4.553 | 0.2076 | 0.15 | 0.85 | Yes | 0.069 | 0.069 |
| M_Yemen | Mota_EN | NWRussia_HG | 3 | 2.06 | 0.5601 | 0.39 | 0.61 | Yes | 0.058 | 0.058 |
| Kotias_HG | Mota_EN | NWRussia_HG | 3 | 2.689 | 0.4421 | 0.389 | 0.611 | Yes | 0.093 | 0.093 |
| Natufian | Mota_EN | NWRussia_HG | 3 | 3.92 | 0.2702 | 0.231 | 0.769 | Yes | 0.152 | 0.152 |
| Zagros_N | Mota_EN | NWRussia_HG | 3 | 0.223 | 0.9739 | 0.484 | 0.516 | Yes | 0.087 | 0.087 |
| Satsurblia_HG | Mota_EN | NWRussia_HG | 3 | 1.595 | 0.6605 | 0.399 | 0.601 | Yes | 0.106 | 0.106 |
| Iberomaurusian | Mota_EN | NWRussia_HG | 3 | 11.054 | 0.0114 | 0.602 | 0.398 | Yes | 0.104 | 0.104 |
| Cluster_F | Mota_EN | NWRussia_HG | 3 | 2.883 | 0.4100 | 0.349 | 0.651 | Yes | 0.061 | 0.061 |
| Cluster_A | Mota_EN | NWRussia_HG | 3 | 3.177 | 0.3651 | 0.491 | 0.509 | Yes | 0.055 | 0.055 |
| Cluster_D | Mota_EN | NWRussia_HG | 3 | 17.712 | 0.0005 | 0.454 | 0.546 | Yes | 0.081 | 0.081 |
